# Reconstruction of 1,000 projection neurons reveals new cell types and organization of long-range connectivity in the mouse brain

**DOI:** 10.1101/537233

**Authors:** Johan Winnubst, Erhan Bas, Tiago A. Ferreira, Zhuhao Wu, Michael N. Economo, Patrick Edson, Ben J. Arthur, Christopher Bruns, Konrad Rokicki, David Schauder, Donald J. Olbris, Sean D. Murphy, David G. Ackerman, Cameron Arshadi, Perry Baldwin, Regina Blake, Ahmad Elsayed, Mashtura Hasan, Daniel Ramirez, Bruno Dos Santos, Monet Weldon, Amina Zafar, Joshua T. Dudmann, Charles R. Gerfen, Adam W. Hantman, Wyatt Korff, Scott M. Sternson, Nelson Spruston, Karel Svoboda, Jayaram Chandrashekar

**Affiliations:** Janelia Research Campus, Howard Hughes Medical Institute, Ashburn, VA 20147, USA; Amazon Web Services, Seattle, WA 98101, USA; Laboratory of Molecular Genetics, The Rockefeller University, New York, NY 10065, USA; Leap Scientific LLC, Hooksett, NH 03106, USA; Environmental Systems Research Institute, Redlands, CA 92373, USA; Intramural Research Program, National Institute of Mental Health, Bethesda, MD 20892, USA

## Abstract

Neuronal cell types are the nodes of neural circuits that determine the flow of information within the brain. Neuronal morphology, especially the shape of the axonal arbor, provides an essential descriptor of cell type and reveals how individual neurons route their output across the brain. Despite the importance of morphology, few projection neurons in the mouse brain have been reconstructed in their entirety. Here we present a robust and efficient platform for imaging and reconstructing complete neuronal morphologies, including axonal arbors that span substantial portions of the brain. We used this platform to reconstruct more than 1,000 projection neurons in the motor cortex, thalamus, subiculum, and hypothalamus. Together, the reconstructed neurons comprise more than 75 meters of axonal length and are available in a searchable online database. Axonal shapes revealed previously unknown subtypes of projection neurons and suggest organizational principles of long-range connectivity.

## Introduction

Mammalian neurons possess extensive axonal arbors that project over long distances (Anderson et al., 2002; Braitenberg and Schüz, 2013; Kita and Kita, 2012; Kuramoto et al., 2013; Wu et al., 2014). These projections dictate how information flows across brain areas. Interareal connections have been studied using tracers that label populations of neurons (Gerfen and Sawchenko; Hunnicutt et al., 2014; Luppi et al., 1990; Markov et al., 2014; Oh et al., 2014; Veenman et al., 1992; Zingg et al., 2014) or with functional mapping methods at various spatial scales (Greicius et al., 2009; Petreanu et al., 2007). However, these methods average across large groups of neurons, including multiple cell types, and obscure fine-scale spatial organization. Mapping brain-wide connectivity at the single neuron level is crucial for delineating cell types and understanding the routing of information flow across brain areas, but very few complete morphological reconstructions of individual neurons are available, especially for long-range projection neurons (Ascoli and Wheeler, 2016; Svoboda, 2011).

Morphological reconstruction is technically challenging because thin axons (diameter, ~100 nm; Anderson et al., 2002; De Paola et al., 2006; Shepherd and Harris, 1998) travel over long distances (centimeters) and across multiple brain regions (Economo et al., 2018; Kita and Kita, 2012; Oh et al., 2014). High-contrast, high-resolution, brain-wide imaging is therefore required to detect and trace axons in their entirety. Earlier studies using imaging in serial sections have reconstructed only small numbers of cells and mostly only partially, because of the difficulty of manual tracing across sections (Blasdel and Lund, 1983; Cowan and Wilson, 1994; Ghosh et al., 2011, 2011; Igarashi et al., 2012; Kawaguchi et al., 1990; Kisvárday et al., 1994; Kita and Kita, 2012; Kuramoto et al., 2009, 2015; Oberlaender et al., 2011; Ohno et al., 2012; Parent and Parent, 2006; Ropireddy et al., 2011; Wittner et al., 2007; Wu et al., 2014). Methods in which a precisely assembled volume is generated, for example based on block-face imaging, are more efficient for tracing (Economo et al., 2016; Gong et al., 2016; Han et al., 2018; Lin et al., 2018; Portera-Cailliau et al., 2005), but manual reconstruction has remained a limiting factor. An RNA sequencing-based method (MAPSeq; Han et al., 2018; Kebschull et al., 2016) has provided a complementary, higher throughput approach to examine single neuron projections. However, this method has two significant limitations: the inherent sensitivity of MAPSeq is unknown and the spatial resolution is lower by orders of magnitude – limited by the volume of tissue that can be micro-dissected for sequencing. Microscopy-based neuronal reconstructions therefore remain the ‘Gold Standard’ for analysis of connectivity and spatial organization of axonal projections.

We previously described a serial 2-photon tomography system for imaging the entire brain at sub-micrometer resolution, and manually tracing fine-scale axonal processes across the entire brain (Economo et al., 2016). Here, we improved this method and developed a semi-automated, high-throughput reconstruction pipeline to efficiently trace many neurons per brain. We completely reconstructed more than 1,000 neurons in the neocortex, hippocampus, thalamus and hypothalamus. Reconstructions were made available in an online database with extensive visualization and query capabilities (www.mouselight.janelia.org). We uncovered new cell types and found novel organizational principles governing the connections between brain regions.

## Results

### High-resolution and high-contrast imaging of the mouse brain

We sparsely labeled neuronal populations by injecting a mixture of low-titer AAV Syn-iCre and a high-titer Cre-dependent reporter (eGFP or tDTomato; Figure 1A), resulting in high fluorescent protein expression in approximately 20-30 cells per injection site (Economo et al., 2016; Xu et al., 2012). Whole brains were harvested, optically cleared, and imaged with an automated two-photon block-face imaging microscope at diffraction-limited resolution and near-Nyquist sampling (voxel size, 0.3 x 0.3 x 1.0 μm^3^; Figure 1B). The entire brain was imaged as a series of partially overlapping image stacks (approximately 40,000 stacks per brain across two channels, 1024 x 1536 x 250 voxels per stack, 5 trillion voxels; imaging time, approximately one week; Figure 1C and 1D; see Supplementary Methods). Image stacks were stitched into a single coherent volume with a non-rigid transform calculated from matching features within overlapping regions (Figure S1A-E). Stitching was accurate at the sub-micrometer level such that small axonal processes were contiguous across stack boundaries and sections (Figure 1E-G).

**Figure 1.**
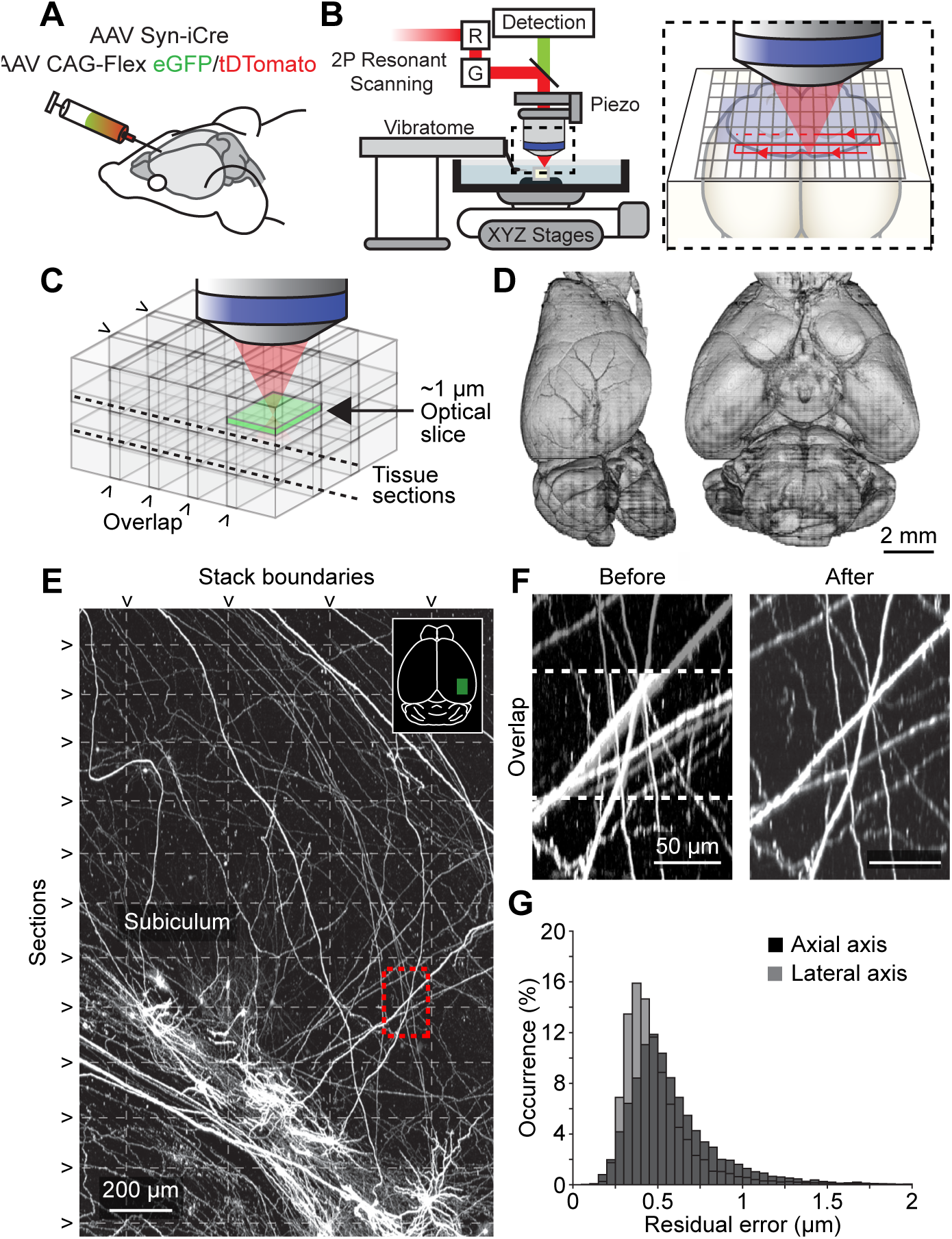
Imaging pipeline. (**A**) Animals were injected in targeted brain areas with a combination of a low-titer AAV Syn-iCre and a high-titer AAV CAG-Flex-(eGFP/tDTomato). (**B**)Two-photon microscope with an integrated vibratome. Inset, sequential imaging of partially overlapping image stacks. (**C**) Image stacks overlapped in x, y, and z. (**D**) Rendered brain volume after stitching (sample 2016-10-31 in Table S2). (**E**) Horizontal maximum intensity projection through a 1300 x 2000 x 600 μm^3^ volume of the motor cortex containing labelled somata and neurites. Horizontal dashed lines mark physical tissue sections; vertical dashed line represents stack boundaries. Dashed box is region shown in F. (**F**) Example of boundary region between two adjacent image stacks before (left) and after stitching (right). Dashed line indicates overlap region. (**G**) Residual stitching error in the lateral and axial directions.

To enhance the throughput of our pipeline we increased the number of cells that could be reconstructed per imaged brain. First, by injecting up to five separate areas per brain we labelled larger numbers of cells, while retaining sparse labeling in any one brain region (Table S1). The fluorescent reporter for each injection site was chosen to minimize the overlap of axons of the same color (e.g. eGFP label for thalamus and tdTomato for the thalamus-projection zones in cortex). Next, we employed a whole-mount immunohistochemical labeling method to amplify the fluorescent signal. This improved the signal-to-noise ratio, especially for neurons that appeared dim with native fluorescence, and thus allowed for a larger portion of labelled neurons to be reconstructed (Table S2). Finally, to ensure optimal image quality throughout the entire sample, we developed a web service that monitored the data acquisition of the microscope (Figure S1F-H). Taken together these improvements resulted in a progressive increase in the number of neurons that could be reconstructed per imaged brain (most recent samples, > 110 reconstructed neurons, n = 3; Table S2).

### Semi-automated reconstruction of complete neurons

Manual interventions during reconstruction remain a significant bottleneck in generating neuronal reconstructions (Economo et al., 2016). Even with specialized custom reconstruction software, a complex cortical neuron previously took approximately 1-3 weeks to be reconstructed in its entirety (Economo et al., 2016). To accelerate reconstruction, we developed a semi-automated reconstruction pipeline consisting of several components. A key first step in the reconstruction workflow is automated segmentation. A classifier trained to identify neurites, including axonal and dendritic processes, was applied to the imaged stacks. The derived probability map was thresholded, skeletonized, and fitted with line segments (Figure 2A and 2B; up to 100K segments per brain). Because crossing points were sometimes misidentified as branch points by the automated segmentation, segments were “broken” at all branch points and close crossing points, to be reconnected later using manual linking. This approach provided more accurate reconstructions compared to full automation. Generated segments covered the majority of axonal length (93% of total length, based on 27 neurons), keeping manual tracing to a minimum.

**Figure 2.**
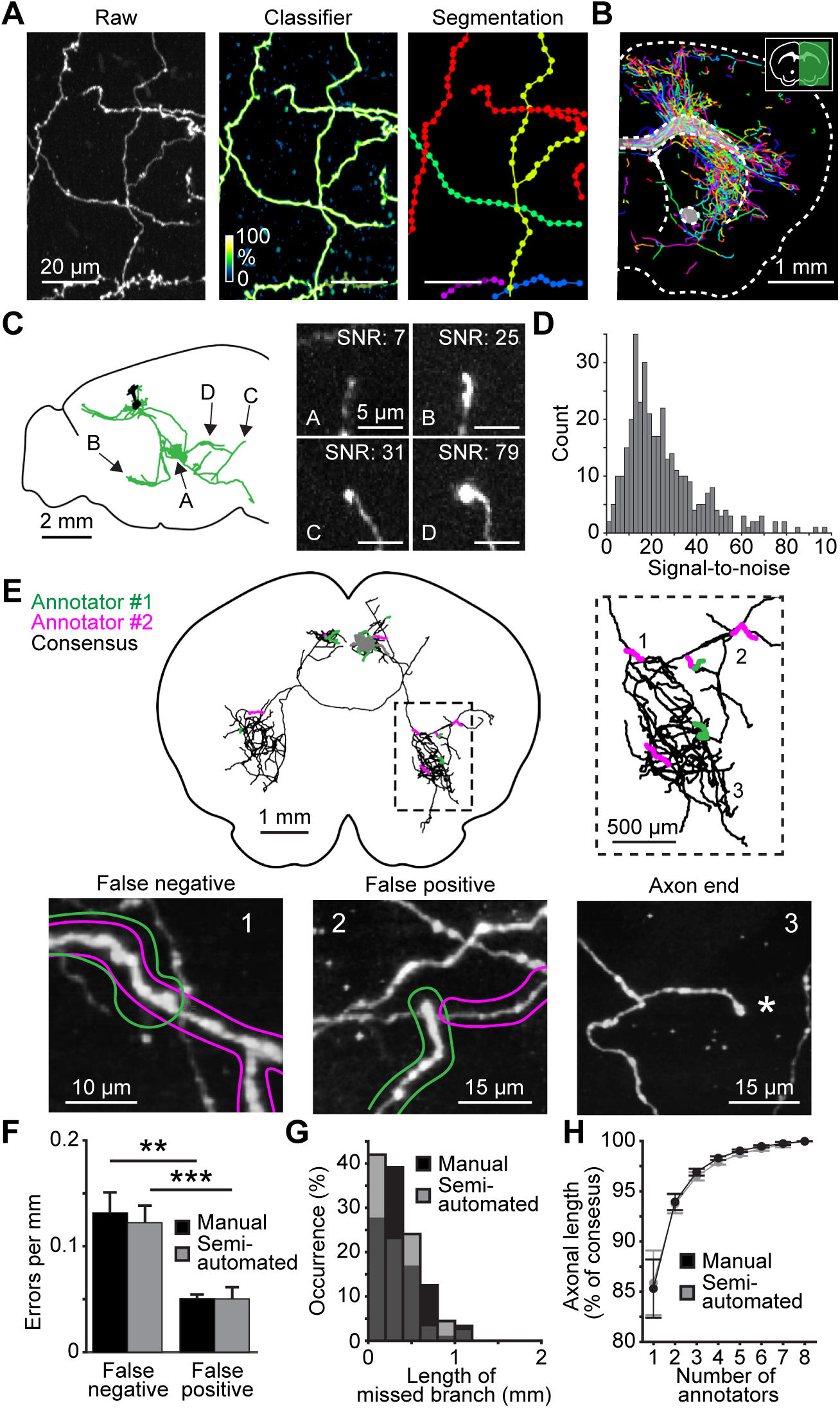
Complete semi-automated reconstructions of individual neurons. **(A)** Automated segmentation. Left, raw maximum intensity image of axons in the thalamus. Middle, probability map output from classifier. Right, segmentation result, with each detected segment depicted in a different color. (**B**) Coronal section through a sample with overlaid segmented neurites. (**C**) Sagittal view of a reconstructed cortical neuron. Arrows indicate location of axon endings several millimeters away from the soma (A, 4.1 mm; B, 4.9 mm; C, 6.5 mm; D, 4.6 mm). (**D**) Distribution of image signal-to-noise ratio for axon endings (difference in intensity between the axon ending and surrounding background, divided by the standard deviation of background). (**E**) Establishing consensus in reconstructions produced by two annotators. Top, example reconstruction of a single neuron. Black, agreement between two annotators; green and magenta, unique segments detected by one of the two annotators. Inset, higher magnification view of dashed box on the left. Bottom, images of the areas of confusion numbered in the inset shown above. Colored outlines represent the reconstructions of the two annotators. (1) False-negative error, or a missed branch. (2) False-positive error consisting of an erroneously appended branch. (3) Axon end correctly identified by both annotators. (**F**) Frequency of each error-type for manual and semi-automated reconstructions. (**G**) Length distribution of missed neuronal branches for manual and semi-automated reconstructions. (**H**) Increase in accuracy as a function of the number of annotators reaching consensus. Accuracy is the proportion of axonal length that was correctly reconstructed. The consensus reconstruction from all eight annotators was defined as 100% accurate.

For manual linking and proofreading, we developed software for efficient three-dimensional visualization and annotation (Janelia Workstation; Murphy et al., 2014), which allowed seamless exploration of the entire mouse brain at multiple resolutions with the overlaid auto-generated line segments. Human annotators used the Workstation to complete the reconstruction by proofreading the segmentation and linking appropriate segments (see Supplementary Methods). Multiple annotators worked collaboratively on the same brain volume concurrently. This semi-automated pipeline resulted in a more than five-fold increase in reconstruction speed compared to manual reconstruction, without any loss in accuracy (manual: 4.5 ± 2.8 mm/hr, Economo et al., 2016; semi-automated: 25.2 ± 11.9 mm/hr). A typical cortical neuron with 10 centimeters of axon was completed in approximately 4 hours and a complex CA3 neuron of 27 centimeters in 8 hours (Figure S2).

Inaccurate reconstructions can arise from poor signal-to-noise ratio or other defects in the data. For this reason, we selectively reconstructed neurons with high and consistent fluorescent signal throughout the entire axonal arbor (Table S2). In these neurons axonal endpoints were clearly identifiable, even at locations far from the soma (Figure 2C and 2D and Figure S3). Neurons that did not meet these criteria were abandoned.

Reconstruction errors can also arise from occasional attentional drift of individual annotators (Helmstaedter et al., 2011). For example, an annotator may overlook a branch point during proofreading and thus miss part of the axonal arbor. To measure these effects, multiple annotators (n = 8) independently traced the same neuron and compared their reconstructions in pairs (Figure 2E). Reconstructions produced by a single annotator had few errors (1.8 ± 0.7 errors per cm; range 1.0 – 2.7; Figure 2F), were typically accurate to almost 90 percent (85.9 ± 9.1% of total axonal length), and were not degraded by the faster semi-automated reconstruction approach (Figure 2F-H). Even more accurate reconstructions were achieved after two annotators compared their work and produced a consensus reconstruction (93.7 ± 4.9 % of total length). Adding a third annotator produced smaller gains in accuracy (96.5 ± 3.1 % of total length; Figure 2H). To balance accuracy and throughput, every neuron in our database was reconstructed by two annotators.

### Online database of more than 1,000 reconstructed neurons

The approach described above allowed us to accurately reconstruct the complete neuronal morphology of more than one thousand neurons. Reconstructed neurons were registered to the Allen Reference Atlas CCF v3 (Gilbert and Ng, 2018) using a landmark registration approach (Figure S4; see Supplementary Methods). More than 1,000 fully reconstructed and registered neurons were deposited in an online resource (www.mouselight.janelia.org). The web interface allows neurons to be searched based on Boolean logic using somatic and axonal target locations (Figures 3-5). Neuronal reconstructions can be viewed together with anatomical structures in 3D and also downloaded for offline analysis. Currently, the database contains projection neurons primarily from the hypothalamus (Figure 3), hippocampus (Figure 4), cerebral cortex (Figures 5-6), and thalamus (Figure 7 and S5). Together the projections from these structures innervate the majority of the brain and are characterized by distinct innervation patterns.

**Figure 3.**
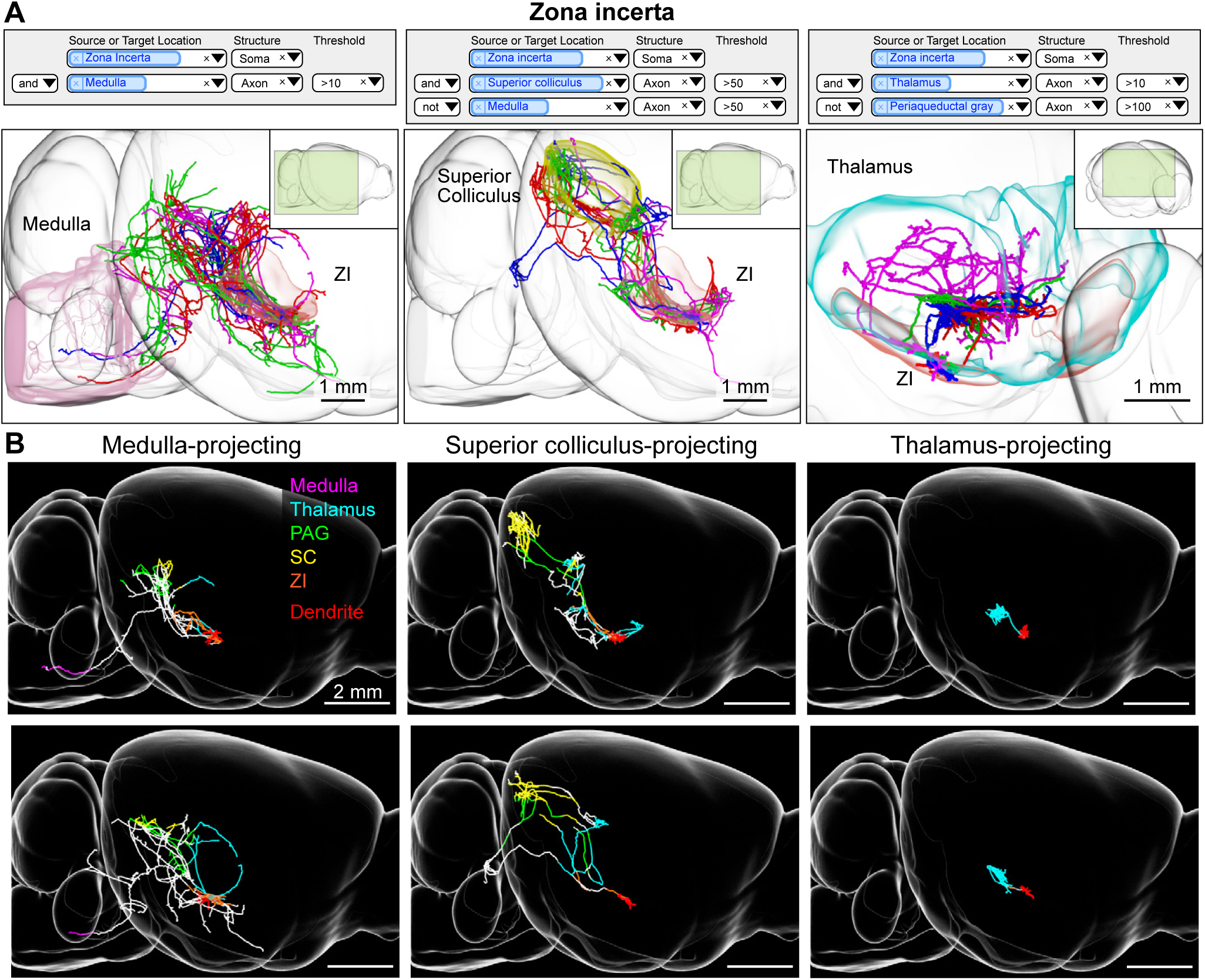
Zona incerta neurons. (**A**) Example queries (top) and 3d visualizations (bottom) for three projection groups in the zona incerta (ZI). Left, ZI neurons with axonal projections in the medulla. Middle, ZI neurons with projections in the superior colliculus but not the medulla. Right, ZI neurons with projections in the thalamus and not the periaqueductal gray. Inset shows perspective of shown area relative to the entire brain. (**B**) Examples of single ZI neurons belonging to the projection groups shown in A. Axons are color coded according to anatomical position (PAG: periaqueductal gray, SC: superior colliculus, ZI: zona incerta). Dendrites are shown in red.

**Figure 4.**
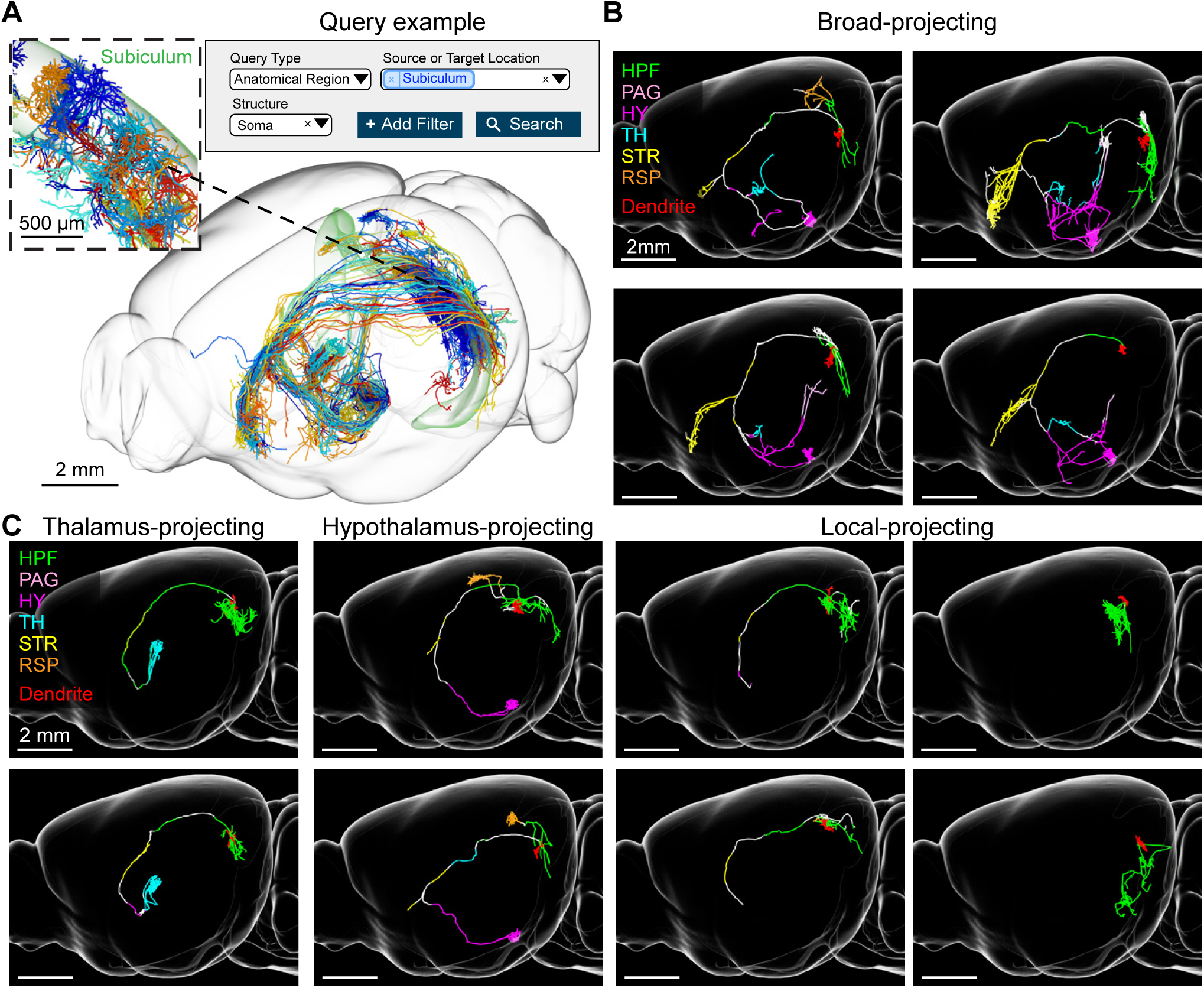
Subiculum neurons. (**A**) Example query requesting all neurons with somata in the subiculum (top) and the resulting 3d visualization (bottom). Inset, higher magnification view of area in dashed box showing the dendrites of the same cells (**B-C**) Example single neuron reconstructions of subiculum neurons with their axons color coded by anatomical position (HPF: hippocampal formation, PAG: periaqueductal gray, HY: hypothalamus, TH: thalamus, STR: striatum, RSP: retrosplenial cortex). Dendrites are shown in red. (**B**) Broad-projecting neurons. (**C**) Thalamus, hypothalamus, and local-projecting neurons.

**Figure 5.**
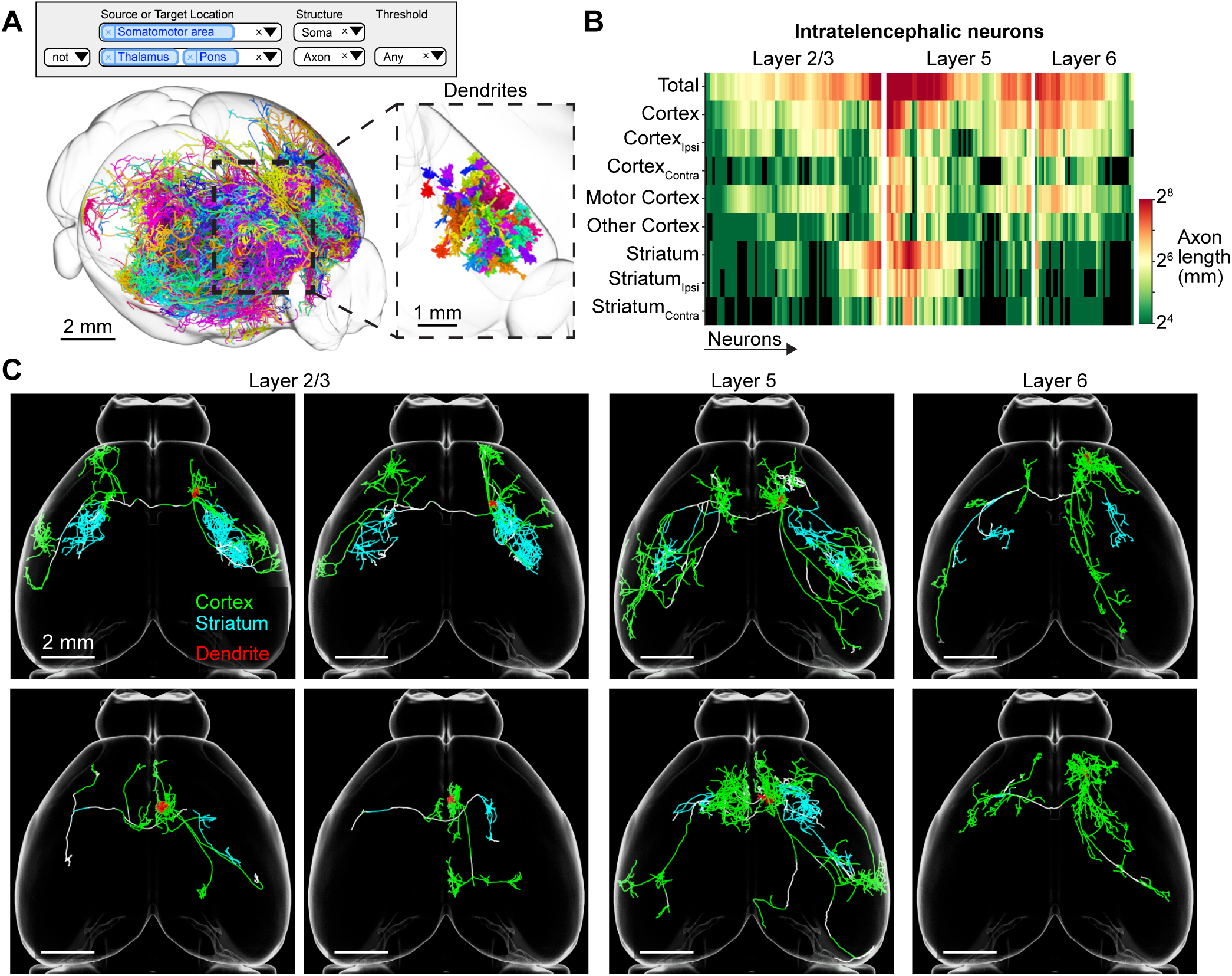
Motor cortex intratelencephalic neurons. (**A**) Search query (top) and 3d visualization (bottom) for intratelencephalic (IT) neurons in the motor cortex classified by lack of axons in thalamus or pons. Inset, higher magnification view of area in dashed box showing the dendrites of the same cells. (**B**) Innervation of telencephalic targets by IT neurons. Rows correspond to projection targets. Columns represent individual IT neurons. Color denotes the axonal length for that cell in a specific area. (**C**) Horizontal view of individual IT neurons with axons color coded according to their anatomical position. Dendrites are shown in red.

With this resource we were able to discover novel projection classes in diverse brain areas. For example, zona incerta (ZI) is a brain region with poorly defined circuits that have been associated with defensive behavior (Chou et al., 2018), appetite (Zhang and van den Pol, 2017), and motor functions (Kim et al., 1992). ZI neurons project to numerous regions within the medulla, midbrain, and thalamus (Sita et al., 2007), but little is known about whether they form projection classes to subsets of these brain regions. We found that rostral ZI neurons are anatomically diverse but could be divided into cells that projected to the periaqueductal grey (PAG) and medulla (Figure 3; n = 4/12 cells), PAG and superior colliculus (SC; n = 4/12), or thalamus (4/12 cells). Two of these cell groups share a projection to the PAG, which is associated with defense and escape behaviors (Chou et al., 2018). PAG and medulla projecting neurons had dense projections to the pontine reticular nucleus, which is involved in movement generation, whereas the other PAG projecting subset innervated SC, a brain area thought to be critical for sensory-guided orientation of movements (Kim, Gregory et al. 1992). Thus, discrete groups of ZI projection patterns appear to map onto different behavioral functions.

Similarly, we discovered clearly delineated projection groups in the subiculum, the main output structure of the hippocampus. This area has been extensively studied for its role in memory, navigation, and motivated behavior (Aggleton and Christiansen, 2015). Neurons in the subiculum formed four groups (Figure 4 and Figure S6-9). One group consists of neurons that have only local collaterals (n = 6), or one unbranched long-range axon (n = 9; Figure S6). Two groups are distinguished by a long-range projection with extensive arborization in either thalamus (n = 13; Figure S7) or hypothalamus (n = 14; Figure S8), with additional projections to other areas. The fourth group (broad-projecting) has multiple targets including thalamus, hypothalamus, periaqueductal gray, and striatum (n = 9; Figure S9). Previous studies based on labeling with pairs of retrograde tracers have suggested that distinct subiculum neurons project exclusively to one long-range target (Naber and Witter, 1998). However, our data set reveals that, only half of the subiculum neurons follow this rule, whereas the other half innervate two or more long-range targets (see also Cembrowski et al., 2018a). Because subiculum neurons project to half a dozen areas in various combinations, the use of retrograde tracers in pairwise combinations have in the past inefficiently detected these complex projection patterns (Kim and Spruston, 2012; Naber and Witter, 1998).

Projection neurons in the cerebral cortex are known to have complex axonal arborizations (Binzegger et al., 2004; Economo et al., 2016, 2018; Han et al., 2018; Kita and Kita, 2012; Oberlaender et al., 2011; Shepherd, 2013). Our database contains motor cortex neurons corresponding to the major cortical projection classes: (1) intratelencephalic neurons (IT) in layers 2-6 (n = 189), (2) pyramidal tract neurons (PT) in layer 5b (n = 32), and (3) corticothalamic neurons (CT) in layer 6 (n = 73; Shepherd, 2013). Based on gene expression analysis, each of these projection classes is thought to contain many subtypes (Arlotta et al., 2005; Economo et al., 2018; Tasic et al., 2018). To date, the diversity in their axonal projections has not been systematically explored.

In our database, IT neurons were some of the most diverse and morphologically complex cells (Figure 5A and 5B). Projections from these neurons are mostly limited to the cortex and striatum, with more minor projections to the basolateral amygdala and the claustrum (Figure 5 and S10; Shepherd, 2013). Collectively, IT neurons projected to (from strongest to weakest projection) motor cortex, somatosensory cortex, insula, ectorhinal cortex, piriform cortex, visual cortex, claustrum, basolateral amygdala. The majority of IT neurons crossed the corpus callosum to project bilaterally (n = 147/175; Figure S10A). The axonal arbors of bilaterally projecting neurons were often strikingly mirror-symmetric across the midline (Yorke and Caviness, 1975), more so for L5 neurons than the others (p < 0.001, one-way ANOVA; L5: 0.36 ± 0.12 Jaccard similarity coefficient, L2/3: 0.26 ± 0.14,, L6: 0.28 ± 0.09; see Supplementary Methods). The majority of IT neurons also projected to the striatum (n = 148/175), either bilaterally (n = 97/175), ipsilaterally (n = 37/175), or contralaterally (n = 14/175).

Cortical projection neurons are often referred to by the efferent projection target of interest, typically identified by retrograde labeling or antidromic activation (e.g., ‘callosal projecting neurons’ or ‘corticostriatal neurons’; Fame et al., 2011; Shepherd, 2013; Turner and DeLong, 2000). In our sample, the majority of IT neurons project both to the striatum and across the corpus callosum. The proportion of axonal length in the striatum or cortex varied by more than one order of magnitude, and was independent of the size of the axonal arbor (Figure S10B). In general, these neurons did not fall into discrete clusters and defied traditional classifications, projecting instead to multiple targets in almost all possible combinations. For example, individual IT neurons can have 40 cm of axon and project to many distinct cortical areas and the striatum; other IT neurons project to a small subset of these areas (Figure 5C and S10C). The diversity in possible projection patterns is much greater than those observed in the subiculum or zona incerta.

### Parallel projections from motor cortex to the thalamus

Diversity in axonal projection patterns give rise to parallel streams of information between connected brain areas. Our database contains numerous reconstructions from the motor cortex and thalamus (Figure 5-7 and S10-13), areas that are bidirectionally interconnected (Guo et al., 2017). Although the reciprocal connections between these areas have been studied (Deschênes et al., 1994; Hunnicutt et al., 2014; Oh et al., 2014), relatively little is known about the projections of individual cells.

Cortical projections to the thalamus originate from layers 5 and 6 (Sherman, 2016). In layer 5 these projections arise from pyramidal tract (PT) neurons, which define layer 5b, and project to the midbrain, brainstem and spinal cord. Their axons do not cross the corpus callosum to the contralateral hemisphere and have limited projections within cortex (Figure 6A; Shepherd, 2013). Corticothalamic projections from layer 5 originate from a genetically distinct subtype of PT neuron (PT thalamus-projecting) in the upper part of L5b (Figure 6B; Economo et al., 2018). These neurons innervate multiple areas of the thalamus, but lack arborizations in the medulla that are characteristic of PT neurons located in lower L5b (PT medulla-projecting).

**Figure 6.**
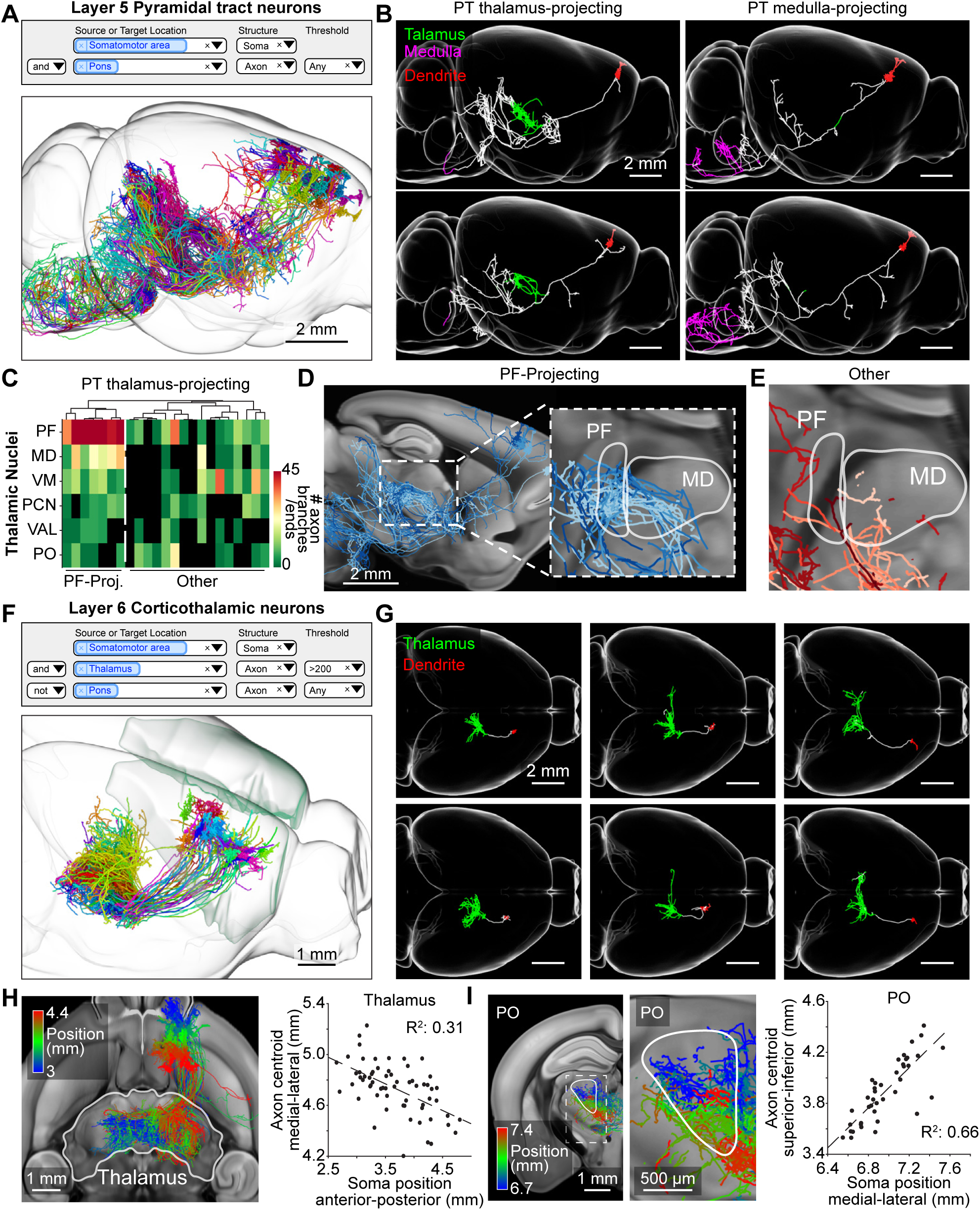
Thalamus-projecting neurons in the motor cortex. (**A**) Search query (top) and 3d visualization (bottom) for pyramidal tract (PT) neurons in the motor cortex classified by the presence of axonal projections in the pons. (**B**) Sagittal view of single PT neurons in upper and lower layer 5, that project to the thalamus (PT thalamus-projecting; green) or medulla (PT medulla-projecting; magenta) respectively. (**C**) Innervation of PT thalamus-projecting neurons to different nuclei of the thalamus. Rows represent key innervated thalamic nuclei, indicated by their acronym (PF: parafascicular nucleus, MD: mediodorsal nucleus, VM: ventral medial nucleus, PCN: paracentral nucleus, VAL: ventral anterior-lateral complex, PO: posterior complex). Each column indicates individual neurons. The color of the heat map denotes the total number of axonal ends and branch points for a neuron in that area. White line represents the separation between neurons with dense innervation in the PF (PF-projecting) and those without (Other). (**D**) Sagittal view of PF-projecting neurons in different shades of blue. Inset, higher magnification view of dashed box on the left (**E**) View of same area shown, with PT thalamus-projecting neurons that do not project to PF in different shades of red. (**F**) Search query (top) and 3d visualization (bottom) of layer 6 CT (L6-CT) neurons in the motor cortex. Neurons are identified by the presence of axons in the thalamus and the lack of projections to the pons. (**G**) Horizontal view of single L6-CT neurons and their axonal projections to the thalamus (green). Dendrites are shown in red. (**H**) Topographic mapping of L6-CT neurons. Left, horizontal section of L6-CT neurons projecting to the thalamus. Individual neurons are color coded according to their somatic anterior-posterior position. Right, relationship between the somatic location of each neuron and the medial-lateral position of its axonal projections within the ipsilateral thalamus. (**I**) Left, coronal section through posterior complex (PO) with axonal projections color coded by their medial lateral position in the motor cortex. Inset, higher magnification view of dashed box on the left. Right, relationship between medial-lateral soma position and superior-inferior axonal position in PO. Trend lines depict linear fit. Axonal position of individual neurons is determined by the weighted centroid of their axonal projections. Positions are in CCF coordinates. Greyscale images are from the Allen reference atlas.

Here we further subdivided the axonal ramifications of PT thalamus-projecting neurons and found that each neuron targets a subset of thalamic nuclei (Figure 6C-E). One subgroup of neurons (n = 7/23 cells) had extensive axonal ramifications in the parafascicular (PF) and mediodorsal (MD) nucleus of the thalamus (Figure 6C). These cells also projected to common extra-thalamic targets: the external segment of the globus pallidus (GPe) and the nucleus of the posterior commissure (NPC; Figure S11). In the cortex they were interspersed with PT thalamus-projecting neurons that project to other parts of the thalamus, such as the ventral medial nucleus (VM), paracentral nucleus (PCN), and the posterior complex (PO) and did not project to GPe or NPC (Figure 6C-E and Figure S11). These findings show that thalamus-projecting PT cells in the motor cortex fall into subtypes that have distinct targets. Additional studies are required to determine how these projection-types correspond to molecular subtypes.

We next investigated the organization of corticothalamic axons originating from layer 6 of the motor cortex (L6-CT). L6-CT cells project almost exclusively to the thalamus, apart from local collaterals in the cortex, and do not cross the corpus callosum (Figure 6F; n = 63 cells; Thomson, 2010). Compared to layer 5 PT thalamus-projecting neurons, L6-CT projections extended over a larger thalamic area (Figure 6G; L6-CT: area = 1.2 ± 0.4 mm^3^, p<0.001, length = 40.8 ± 12.5 mm, p<0.001, PT thalamus-projecting: area = 0.7 ± 0.4 mm^3^, length = 16.9 ± 10.7 mm; see Supplementary Methods) and exhibited about twice as much branching of axonal arbors within the thalamus (L6-CT = 80.3 ± 49.9 branch points, PT thalamus-projecting = 34.9 ± 26.7 branch points; p<0.001). The L6-CT neurons projected to diverse thalamic nuclei. Most neurons had axonal ramification in either PO, MD, or reticular nucleus (RT; Figure S12). However, these projections were not always mutually exclusive. Whether L6-CT neurons in the motor cortex form discrete cell-types, as has been suggested in the sensory cortex (Hoerder-Suabedissen et al., 2018; Shima et al., 2016), or represent a continuum of projection patterns will require additional investigation.

L6-CT neurons in the somatosensory cortex project to sensory thalamus in a topographically organized manner (Deschênes et al., 1998). Our single-neuron reconstructions reveal a similar topographic organization in the motor cortex. L6-CT neurons in the posterior motor cortex sent their axonal projections to lateral regions of the thalamus (Figure 6H; R2 = 0.31, p < 0.001 linear regression). Furthermore, these posterior neurons mostly remained on the ipsilateral side of the thalamus, unlike more anterior neurons (R2 = 0.17, p < 0.001 linear regression; Figure S13A). Topographic maps organized along distinct axes were also seen within targeted thalamic nuclei of L6-CT neurons (Figure 6I and S13B-C).

### VAL thalamus contains fine-scale projection maps

Neurons in the ventral anterior-lateral complex of the thalamus (VAL) projected to a large area of the ipsilateral motor cortex (Bosch-Bouju et al., 2013), with additional projections to the somatosensory cortex (n = 34 cells). Some neurons also arborized in RT (n = 4/34) and the dorsal striatum (n = 14/34; Figure 7A-B; Kuramoto et al., 2009). Individual VAL neurons had arborizations confined to small areas of the cerebral cortex. These projections were not random; instead, groups of neighboring neurons in VAL had similar thalamocortical projections (Figure 7C). For example, neurons within the caudomedial VAL had two main axonal tufts located in the anterior motor and sensory cortex respectively (n = 18/34 cells; Figure 7D). These two branches were positioned around a common plane of symmetry that corresponded to the border between the motor and somatosensory cortical areas. The relative distance of each tuft to the anatomical border correlated with the neurons mediolateral position in VAL (R2 = 0.42, p < 0.01 linear regression; Figure 7E). These findings show fine-scale topography between subregions of VAL and cortical areas of different modalities. Studying the axonal arbors of individual neurons therefore reveals organization of projections that are obscured with traditional bulk labeling approaches.

**Figure 7.**
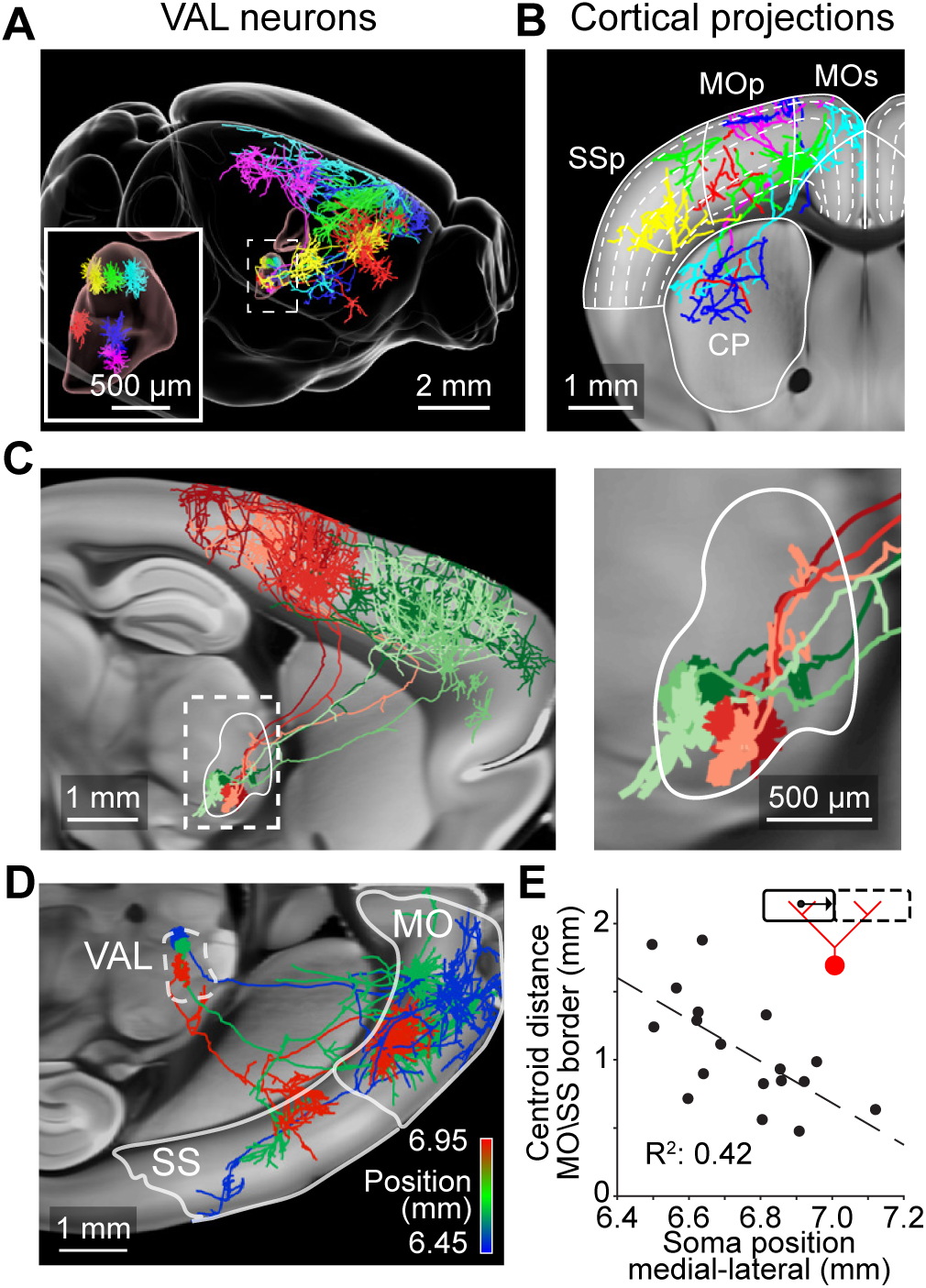
Cortical projections from neurons in the VAL complex of the thalamus. (**A**) Six VAL neurons and their axonal projections. (**B**) Coronal view of the same neurons. Dashed lines show position of cortical layers. (**C**) Left, sagittal view of two groups of VAL neurons with similar projection patterns in the posterior (red; n = 3 cells) or anterior (green; n = 3 cells) motor cortex. Right, higher magnification view of dashed box on the left. (**D**) Horizontal view of cortical projections of three neurons in caudomedial VAL color coded according to their medial lateral position. White outline show location of motor (MO) and sensory cortex (SS). (**E**) Relationship between somatic medial-lateral position and the average distance of the axonal centroid in the motor (MO) and sensory cortex (SS) to the border of those two areas (illustrated by inset). Reported positions are in accordance with the Allen Common Coordinate Framework. Greyscale images are from the Allen reference atlas.

## Discussion

We describe an imaging and reconstruction platform capable of producing complete neural reconstructions at unprecedented speed and scale. More than 1,000 fully reconstructed neurons are now available in our online database (www.mouselight.janelia.org). This database represents a unique resource for the investigation of interareal connectivity and discovery of projection classes in a mammalian brain. For example, clustering based on these data was used to identify putative cell types based on projection patterns (Figures 3-5; see also Economo et al., 2018), as well as putative subtypes within previously described projection classes (Figure 6) and topographic gradients of single-cell projections (Figures 6-7).

Previous large-scale studies of long-range projections have relied on bulk labeling to trace projections across brain areas (Hintiryan et al., 2016; Hooks et al., 2018; Hunnicutt et al., 2014; Oh et al., 2014; Zingg et al., 2014). These approaches effectively average across distinct cell types with idiosyncratic projections, thus potentially obscuring the detailed organization of individual axonal arbors. Neurons with distinct projection patterns have traditionally been identified based on combinatorial retrograde labeling with multiple markers (Betley et al., 2013; Kim and Spruston, 2012; Naber and Witter, 1998; Pan et al., 2010). This approach is cumbersome because the number of required injections scales non-linearly with the number of projection targets. In addition, the spatial specificity is limited by the confinement of the tracer, and biases are produced by variable efficiency of tracer uptake in different brain regions and by different cell types; tracer uptake by axons of passage may also complicate retrograde tracer experiments. These limitations lead to potential false negatives in the detection of combinatorial projections. In contrast, complete neuronal reconstructions provide axonal projection maps at the finest levels of detail and can unambiguously reveal the diversity of multi-areal projection patterns.

Great progress has been made in methods for single-cell RNA sequencing (scRNA-seq), which now provides detailed transcriptomes of individual cells. Large-scale scRNA-seq measurements in the neocortex and other structures have identified known cell types and in addition also discovered additional transcriptomic clusters, which might correspond to newly discovered cell types (Cembrowski et al., 2018b; Shekhar et al., 2016; Tasic et al., 2016, 2018; Zeisel et al., 2018). Establishing the relationship between transcriptomic cluster and cell type requires validation with other types of information, typically neuronal structure. For example, the correspondence between transcriptomic clusters and retinal bipolar cells with distinct morphology is excellent (Shekhar et al., 2016). Pyramidal tract neurons in the motor cortex with distinct transcriptomic clusters correspond to different pyramidal tract neurons with axonal arborizations in either the medulla or thalamus (Economo et al., 2018). Recordings from these cells revealed distinct activity correlated during the planning and execution of goal-directed movements. These studies illustrate how the structural cell types in our database can be linked to molecular cell types and neural function.

In the future, more efficient methods for linking molecular and morphological information will be critical for characterizing neuronal cell types. Since our histological treatments and imaging are non-destructive (i.e. they result in a series of preserved brain slices) our pipeline is amenable to post-hoc gene expression analysis by in-situ hybridization techniques and other types of molecular characterization (Chen et al., 2015; Eng et al., 2017; Shah et al., 2018). The synergy between large-scale genetic (Tasic et al., 2016; Zeisel et al., 2018) and morphological studies is primed to transform our understanding of how the brain is organized.

Single-neuron reconstructions have been performed in the past (Igarashi et al., 2012; Kita and Kita, 2012; Kuramoto et al., 2009; Lin et al., 2018; Wittner et al., 2007; Wu et al., 2014), but at low rates and often not to completion. For example, a single CA3 pyramidal neurons in the hippocampus (Wittner et al., 2007) was reconstructed over four months; a similar CA3 neuron in the MouseLight database was reconstructed within eight hours (Figure S2, length: 27.3 cm). Several advances contributed to the throughput and accuracy required to produce this data set, including improved tissue processing, high-contrast imaging, automated segmentation, and 3d volumetric visualization for proofreading.

The precisely assembled image volume generated by our pipeline is amenable to even faster reconstructions using more advanced computational approaches. Much progress has been made towards fully automated tracing, but accuracy is still limiting (Acciai et al., 2016; Peng et al., 2015). In our pipeline, we segmented neurites in an automated manner, but the segments were linked into trees by manual proofreading. Combining automated segmentation with manual reconstruction allowed us to reconstruct complete neurons, including all termination zones, revealing the full complexity of axonal arbors. A full characterization of the cell types of the mouse brain will likely require reconstructions of 100,000 neurons or more (assuming 1000 brain regions, an average of 10 cell types per brain region, and 10 measurements per cell type; Svoboda, 2011). How can the required increase in throughput be achieved? Despite the advances presented here the speed of reconstructions is still limiting. One brain containing 100 neurons can be imaged in one week per microscope, and the imaging can be accelerated and parallelized. In contrast, with our current semi-automated workflow, reconstructing 100 neurons requires approximately 400 person-hours (10 person-weeks) of manual proofreading. Currently, all decision points (i.e. linking of automatically segmented neurites to a tree) are curated manually, which is the limiting step in reconstructions. We believe that a 100-fold acceleration in reconstruction speed is required and may be feasible if all but the most difficult decisions are performed by computer algorithms in a fully automated manner, for example using convolutional neural networks. Our database of gold-standard reconstructions will serve as training data for machine learning algorithms that will be required to achieve these gains. Together with increased speeds in imaging, this enhanced reconstruction speed would make the goal of a full characterization of the cell types in the mouse brain achievable.

## Supplementary Methods

### Contact for reagent and resource sharing

Further information and requests for resources and reagents should be directed to and will be fulfilled by the Lead Contact, Jayaram Chandrashekar.

### Experimental model and subject details

#### Animals

Wild-type C57BL/6 animals were obtained from Charles River Laboratories. Rorb-IRES2-Cre (IMSR_ JAX:023526; Harris et al., 2014) and Sim1-Cre (IMSR_JAX:006451; Balthasar et al., 2005) transgenic animals were obtained from The Jackson Laboratory. Adult females (>8 weeks) were used for all experiments and were group housed with sex-matched littermates. None of the animals had undergone any previous procedures. Mice were monitored on their health, had access to ad libitum food and water, and were housed in an enriched environment. All experimental protocols were conducted according to the National Institutes of Health guidelines for animal research and were approved by the Institutional Animal Care and Use Committee at Howard Hughes Medical Institute, Janelia Research Campus (Protocol #14–115).

### Method details

#### Viral labelling

To achieve sparse labelling, we injected adult C57/BL6 mice with a combination of highly diluted adeno associated virus expressing Cre-recombinase (AAV Syn-iCre) and high-titer reporter virus coding for a fluorescent reporter (AAV CAG-Flex eGFP/tDTomato). Multiple non-overlapping brain areas were labeled within a single animal (Table S1). For Rorb-IRES2-Cre transgenic animals, sparsity was achieved using diluted Cre-dependent FLP-recombinase (AAV Syn-Flex-FLPo) and a FLP-dependent reporter virus (AAV CAG-FRT-eGFP/tDTomato; Table S2). Sim1-Cre animals received a systemic injection via the retro-orbital sinus with a mixture of Cre-dependent FLP-recombinase (PHP-eB-Syn-Flex-FLPo) and a FLP-dependent reporter virus (PHP-eB-CAG-FRT-3xGFP; Chan et al., 2017). High titer (> 1012 GC/ml) viruses were obtained from the Janelia Research Campus Molecular Biology Core and diluted in sterile water when necessary.

#### Tissue preparation and clearing

Transfected mice were anesthetized with an overdose of isoflurane and then transcardially perfused with a solution of PBS containing 20 U/ml heparin (H3393, Sigma-Aldrich, St. Louis, MO) followed by a 4% paraformaldehyde solution in PBS. Brains were extracted and post-fixed in 4% paraformaldehyde at 4°C overnight (12-14 hours) and washed in PBS to remove all traces of excess fixative (PBS changes at 1h, 6h, 12h, and 1 day).

For imaging of endogenous fluorescence, brains were delipidated by immersion in CUBIC-1 reagent for 3-7 days (Economo et al., 2016; Susaki et al., 2015). For amplification by immuno-labeling, brain were delipidated with a modified Adipo-Clear protocol (Chi et al., 2018). Brains were washed with methanol gradient series (20%, 40%, 60%, 80%, Fisher #A412SK) in B1n buffer (H2O/0.1% Triton X-100/0.3 M glycine, pH 7; 4 mL / brain; one hour / step). Brains were then immersed in 100% methanol for 1 hour, 100% dichloromethane (Sigma #270997) for 1.5 hours, and three times in 100% methanol for 1 hour. Samples were then treated with a reverse methanol gradient series (80%, 60%, 40%, 20%) in B1n buffer for 30 minutes each. All procedures were performed on ice. Samples were washed in B1n buffer for one hour and left overnight at room temperature; and then again washed in PTxwH buffer (PBS/0.1% Triton X-100/0.05% Tween 20/2 µg/ml heparin) with fresh solution after one and two hours and then left overnight.

After delipidation, selected samples were incubated in primary antibody dilutions in PTxwH for 14 days on a shaker (1:1000, anti-GFP, Abcam, #ab290; 1:600, anti-tDTomato, Sicgen, #ab8181). Samples were sequentially washed in 25ml PTxwH for 1, 2, 4, 8, and three times for 24 hours. Samples were incubated in secondary antibody dilutions in PTxwH for 14 days (Alexa Fluor 488 conjugated donkey-anti-rabbit IgG, 1:400, Invitrogen, #A21206; Alexa Fluor 546 conjugated donkey-anti-goat IgG, 1:400, Invitrogen, #A11056) and washed in PTxwH similar to descriptions above.

Brains were embedded in 12% (w/v) gelatin and fixed in 4% paraformaldehyde for 12 hours. Index matching for optical clarity was achieved by immersing the samples in solutions of 40% DMSO in 10 mM PB with increasing concentrations of D-Sorbitol (up to 40/60 w/v; see Economo et al., 2016) or in a solution of 40% OptiPrep (D1556, Sigma-Aldrich) in DMSO. The final imaging medium had a refractive index of 1.468 which allowed for imaging up to depths of 250 μm in the tissue without significant loss of fluorescence (Economo et al., 2016).

#### Microscope

Processed samples were imaged using a resonant scanner two-photon microscope imaging at 16 million voxels per second (Economo et al., 2016). The microscope is integrated with a motorized stage (XY: M-511.DD, Z: M-501-DG, Controller: C843; Physike Instrumente, Karlsruhe, Germany) and vibratome (Leica 1200S, Leica Microsystems, Wetzlar, Germany). Imaging was performed using a 40x/1.3 NA oil-immersion objective (#440752, Carl Zeiss, Oberkochen, Germany) attached to a piezo collar (P-725K.103 and E-665.CR, Physik Instrumente). Image stacks (385×450X250 μm^3^) were collected with a voxel size of 0.3×0.3x 1 μm^3^ in two channels (red and green). The surface of the sample was automatically detected using the difference in autofluorescence between the tissue and the embedding gelatin. Scanning of the exposed brain block-face was then achieved by dividing it into smaller image stacks. Overlap between adjacent stacks (25 μm) allowed feature-based registration and stitching. After the block-face was imaged to a depth of 250 μm the vibratome removed 175 μm of tissue, leaving approximately 75 μm of overlap across imaged sections. The mouse brain was imaged in approximately one week.

To ensure robust and fault-proof processing of our large datasets, we created a custom software pipeline that facilitates the multi-step data processing in real time as the images are being acquired (https://github.com/MouseLightProject). Success or failure of the processing of each tile for each task was tracked, logged, and reported in a graphical user interface (Figure S1H). Each imaged stack was analyzed for possible faults in imaging (e.g. air bubble in front of the objective) or sectioning (e.g. larger than expected slice thickness). In case of a detected fault the microscope was automatically shut down to allow for manual adjustments.

#### Feature-based volume stitching

A fully imaged brain consists of ~20k imaged stacks that need to be stitched to create a coherent volume. Accurate stitching is necessary to eliminate discontinuous neurites at the borders, which is critical for reliable reconstruction and automation. Standard methods involving linear transformations between adjacent tiles, following by global optimization, do not produce micrometer level precision (Bria and Iannello, 2012; Chalfoun et al., 2017; Emmenlauer et al., 2009; Tsai et al., 2011). To account for non-linear deformations (caused by physical sectioning, optical field curvature etc.), we extended the descriptor based stitching framework we described previously (Economo et al., 2016) to all three dimensions. First, blob-like objects were detected in individual tiles using a difference of Gaussian (DoG) filter. These descriptors were matched between adjacent tiles in both x, y, and z directions using a coherent point drift algorithm (Myronenko and Song, 2010). Matched descriptors were then used to estimate a non-rigid transformation which mapped voxel locations to a target coordinate space while preserving the spatial ordering. For computational efficiency the transformation was represented as a set of barycentric transforms of 5×5×4 equally distributed control points in each stack (x, y, z respectively).

#### 3D visualization and reconstruction

For viewing and annotating terabyte-scale image stacks, each dataset (input tiles) was resampled into a common coordinate space according to the transforms determined during the stitching procedure. This produced a set of non-overlapping image stacks (output tiles) that spanned the imaged volume. Resampling was achieved by back-projecting each voxel in each output tile to the nearest sampled voxel. In regions where two or more input tiles overlapped, the maximum intensity was used for the corresponding location in the output tile. The resampling task was implemented on a cluster (64 nodes each with 32 cores and 512 GB RAM) of Intel CPUs with Advanced Vector Instructions 2 (AVX2) and parallel access to high-bandwidth network storage. The input tiles were partitioned into contiguous sets and traversed in Morton order to facilitate merging overlaps in memory. Execution was dominated by the time required to read and write data to disk. Data were resampled to 0.25 × 0.25 × 1 µm voxels, and stored on disk along with downsampled octree representations of the same volume for visualization at different spatial scales.

The imaged data was viewed and annotated in the Janelia Workstation (JW; Murphy et al., 2014) The JW allows low lag, multi-scale visualization of terabyte-scale image volumes, rendering of sub-volumes in three dimensions, and ergonomic annotation tools for tracing and proofreading. A detailed description of the JW will be published elsewhere.

#### Semi-automated segmentation

We trained a binary random forest classifier to identify axonal processes using five different appearance and shape features (Gaussian, Laplacian, gradient magnitude, difference of Gaussian, structure tensor Eigenvalues, Hessian of Gaussian Eigenvalues) at multiple spatial scales (σ = 0.3, 1, 1.6, 3.5, and 5 μm) (Ilastik, Sommer et al., 2011). Training sets were created by pixel-based classification of axonal processes in regions containing different morphological features (axons close to the soma, termination zones etc.). The output of the classifier consisted of a probability stack where the value of each pixel reflects the likelihood of it belonging to an axon. A morphological skeleton (Lee et al., 1994) was then created by extracting the centerline of the classifier output after thresholding (> 0.5). For efficiency during visualization, we downsampled the resulting line segments to approximately 20 μm spacing in between nodes.

All segmented neurites were split into linear segments by separating them at their axonal branch points. Several filtering steps were applied on the resulting segmentation to remove incorrectly identified processes. For instance, we used a path-length based pruning strategy where spurious branches (<15 μm) were deleted. Furthermore, a separate classifier was trained to identify auto-fluorescence originating from fine-scale vasculature and non-specific antibody binding. This approach generated up to 100,000 axonal segments per brain (length after split: 96.5 ± 155.4 μm, before: 263.4 ± 625.6 μm). The coverage of axonal processes by these generated segments was determined using fine-scale reconstructions (internode interval <5 µm) of all axonal projections in the caudoputamen of one sample and manually assigning fragments to each neuron (without merging).

Human annotators started from the soma and reconstructed neurons by linking neurite segments. Annotators in addition filled in parts of the neuron that may have been missed by the segmentation. The location of branch points and axonal endpoints were annotated to ensure that the entire axonal tree was reconstructed. Neurons that could not be reconstructed (with dim neurites or neurites that traversed a densely labeled region) were identified by experienced annotators after approximately 30 minutes and were abandoned (median: 19% of cells in samples with more than 50 cells). Two annotators independently reconstructed each neuron and compared their work to generate a consensus reconstruction, which was stored in the database.

#### Sample registration

Imaged brains were aligned to the Allen Common Coordinate Framework (CCF) by taking a downsampled version of the entire sample volume (~5 x 5 x 15 μm voxel size) and registering it to the averaged Allen reference template (10 x 10 x 10 μm voxel size) using the 3D Slicer software platform (slicer.org, Kikinis et al., 2014). First, parts of the brain that were not present in the reference template (e.g. the anterior olfactory bulb and posterior cerebellum/spinal cord), as well as the imaged gelatin, were cropped out of the sample volume. An initial automated intensity-based affine registration was then performed to align the sample to the reference atlas (BRAINSFit module, Johnson et al., 2007). An iterative landmark-registration processes, using thin-plate splines approximation, was then used to achieve a more precise registration to the CCF (LandmarkRegistration module, 110 ± 60 µm variation between independent registrations of the same sample). All registration steps were used to generate a single displacement vector field which was used to align reconstructed neurons to the CCF.

#### Quantification and statistical analysis

All statistical analysis was performed in Matlab using custom made scripts. Unless stated otherwise, center and dispersion of bar plots represent mean ± SEM and descriptive statistics in the main text represent mean ± std. Significance was defined as p < 0.05, with the specific statistical test provided in main text or within associated figure legend. Sample sizes (n) can refer to the number of samples/ brains, or number of cells and is clarified in the associated text. Significance conventions are as follows: NS: p ≥ 0.05; ٭: p < 0.05; ٭٭: p < 0.01; ٭٭٭: p < 0.001 associated sample sizes are provided in the main text or within the figure legend. All reported position information is according to the Allen CCF.

Axonal length within an anatomical area was measured by taking all nodes within the given area and summing the distances to their parent nodes. To calculate the weighted centroid of axonal projections we resampled the axonal tree graph by linearly sampling at 1 µm interval between connected nodes. Nodes within the area of interest were then selected using 3D meshes from the Allen CCF and their positions were averaged to derive the centroid. To exclude enpassant axons, only neurons with at least three axonal ends were considered. Axonal area size was calculated by dividing the brain area of interest into isotropic voxels (200 x 200 x 200 µm) and counting the number of voxels that an axon passed through. Bilateral symmetry of IT projections in the cortex was measured by similarly dividing the brain into isotropic voxels (1 x 1 x 1 mm) and mirroring one hemisphere onto the other. Each voxel was scored for axon traversal on the ipsilateral and contralateral side. Symmetry was then calculated using the Jaccard similarity coefficient dividing the number of voxels that were positive for both hemispheres with the total number of positive voxels.

#### Data and software availability

All reconstructed neurons are available for download from the MouseLight neuron browser (http://ml-neuronbrowser.janelia.org/). Matlab code for the performed analysis is available online (https://github.com/JaneliaMouseLight/Analysis-Paper).

## Acknowledgements

The authors would like to thank all members of the MouseLight project team; Gordon M. Shepherd, and Giorgio Ascoli for valuable feedback on the manuscript; Nathan Clack for helpful comments and technical assistance; Janelia Experimental Technologies, especially Daniel Flickinger, Vasily Goncharov, and Christopher McRaven, for optical design, construction and support; Janelia Virus Services, especially Kimberley Ritola for viral reagents; Janelia Histology, in particular Monique Copeland, Brenda Shields, and Amy Hu, Janelia Vivarium, especially Salvatore DiLisio, Jared Rouchard, and Sarah Lindo, for animal care and surgical assistance. We would also like to thank Amanda Collins, Najla Masoodpanah, Rinat Rachel Mohar, and Takako Ohashi for additional reconstruction work. We are also grateful to the Allen Institute for Brain Science for providing the Allen Mouse Brain Atlas. This work was supported by the Howard Hughes Medical Institute.

## Author contributions

J.C., K.S., N.S., A.W.H., J.T.D, and S.M.S. conceptualized the study; J.C. managed the project with input from W.K., N.S. and K.S.; J.W., N.S., K.S., and J.C. wrote the manuscript; J.W. conducted the anatomical analysis and made the illustrations, with input from J.C., K.S., N.S., J.T.D., C.R.G., S.M.S., and M.N.E.; J.W., T.A.F., and J.C. prepared the samples and acquired the whole-brain imaging data; Z.W. developed the whole-brain immuno-labelling procedure; E.B. developed the automated segmentation tools and processed the acquired data; E.B. and B.J.A. developed the stitching and rendering software; P.E. developed the online database and the computational pipeline with input from E.B., J.W., and T.A.F.; C.B., D.J.O., K.R., D.S., and S.D.M. developed the data visualization tools; C.B. and D.S. developed the 3D reconstruction software; D.G.A. developed software modules for the automated block-face imaging microscope; C.A., P.B., R.B., A.E., M.H., D.R., B.D.S., M.W., and A.Z. generated the neuronal reconstructions.

---

**Table S1.** Information on viral injections. Related to Figure 1.

| Location | AAV-Cre |  | Coordinates |  |
| --- | --- | --- | --- | --- |
| (abbreviation) | (sereo-type) | (dilution) | (A/P), (M/L) | (depth) |
| MO | 2/1 | 1:15-60k | +2.5, +1.5 | -0.9, -0.4 |
|  |  |  | +1.5, +1.5 | -0.9, -0.4 |
|  |  |  | +1.5, +2.5 | -0.9, -0.6, -0.3 |
|  |  |  | +2.3, +1.2 | -0.9, -0.4 |
|  |  |  | +1.4, +0.8 | -0.8, -0.3 |
| SUBd | 2/1 | 1:30-60k | -3.6, +2.5 | -1.5 |
|  |  |  | -3.05, +1.5 | -1.5, -1.4 |
|  |  |  | -3.8, +2.5 | -1.85 |
|  |  |  | -3.2, +1.6 | -1.6 |
| ANT | 2/5 | 1:20-45k | -0.6, -0.8 | -2.4 |
|  |  |  | -0.7, -0.6 | -3.2, -2.7 |
| VAL | 2/5 | 1:20-45k | -1.8, -0.6 | -4.1 |
|  |  |  | -0.95, -1.2 | -3.4, -2.9 |
|  |  |  | +1.4, -0.8 | -4.0, -3.5 |
| PVH | 2/5 | 1:20-45k | -0.75, -0.3 | -5.0 |
| PLH | 2/5 | 1:20-45k | -0.95, -1.1 | -4.4 |

**Table S2.** Information on imaged brains. Related to Figure 1.

| Brains | Delipidation | RI matching | IHC | Viral injections | Reconstructed neurons |
| --- | --- | --- | --- | --- | --- |
| 2014-06-24 | Cubic R1 | D-Sorbitol | No | 3 | 48 |
| 2015-06-19 | Cubic R1 | D-Sorbitol | No | 2 | 15 |
| 2015-07-11 | Cubic R1 | D-Sorbitol | No | 2 | 25 |
| 2016-04-04 | Cubic R1 | D-Sorbitol | No | 4 | 17 |
| 2016-07-18 | Cubic R1 | D-Sorbitol | No | 2 | 13 |
| 2016-10-25 | Cubic R1 | D-Sorbitol | No | 4 | 23 |
| 2016-10-31 | Cubic R1 | D-Sorbitol | No | 5 | 31 |
| 2017-01-15 | Cubic R1 | D-Sorbitol | No | 5 | 19 |
| 2017-02-22 | Cubic R1 | D-Sorbitol | No | 6 | 14 |
| 2017-04-19 | Cubic R1 | D-Sorbitol | No | 6 | 22 |
| 2017-05-04 | Cubic R1 | D-Sorbitol | No | 5 | 31 |
| 2017-06-10 | Cubic R1 | D-Sorbitol | Yes | 6 | 19 |
| 2017-06-28 | Cubic R1 | D-Sorbitol | No | 6 | 48 |
| 2017-08-10 | Cubic R1 | D-Sorbitol | No | 2* | 11 |
| 2017-09-11 | Adipo-Clear | D-Sorbitol | Yes | 5 | 24 |
| 2017-09-19 | Adipo-Clear | D-Sorbitol | Yes | 5 | 40 |
| 2017-09-25 | Adipo-Clear | D-Sorbitol | Yes | 5 | 55 |
| 2017-11-17 | Adipo-Clear | OptiPrep | Yes | 5 | 62 |
| 2018-03-09 | Adipo-Clear | OptiPrep | Yes | 5 | 57 |
| 2018-04-03 | Adipo-Clear | OptiPrep | Yes | 5 | 7 |
| 2018-04-13 | Adipo-Clear | OptiPrep | Yes | 5 | 144 |
| 2018-05-23 | Adipo-Clear | OptiPrep | Yes | 5 | 24 |
| 2018-06-14 | Adipo-Clear | OptiPrep | Yes | 5 | 14 |
| 2018-07-02 | Adipo-Clear | OptiPrep | Yes | 5 | 154 |
| 2018-08-01 | Adipo-Clear | OptiPrep | Yes | 1** | 115 |

\* Rorb-Cre-1 animal.
\*\* Sim1-Cre animal.

**Figure S1.**
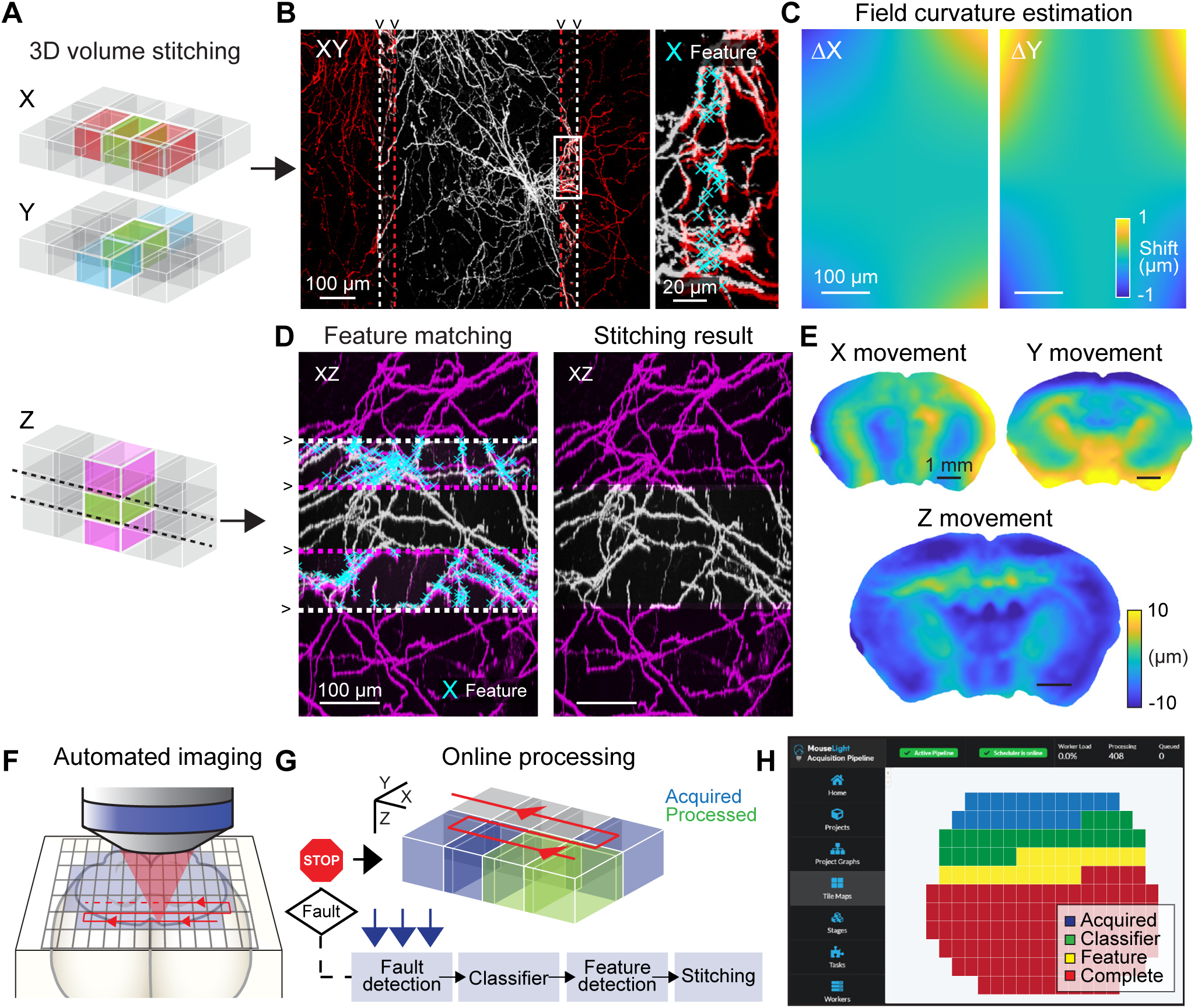
Two-photon serial tomography pipeline. Related to Figure 1. (**A**) Schematic representation of 3D volume stitching approach. Individual stacks are registered based on matching features with neighboring stacks in all three dimensions. (**B**) Coronal maximum projection of one example imaged stack with two neighboring stacks shown in red. Dashed lines indicate the stack boundaries that mark the overlap region between two stacks. Inset shows expanded view of the overlap region with detected image features marked as cyan crosses. (**C**) Example of the estimated error between neighboring tiles due to optical field curvature distortion in the x and y direction. (**D**) Left, neighboring image stacks in the axial plane. Right, result of the complete stitching process. (**E**) Example of the estimated movement after sectioning in all three dimensions. (**F**) Sequential imaging of overlapping image stacks. Red line indicates the direction of imaging. (**G**) Schematic of online processing pipeline. Imaged stacks are automatically processed as they become available. Acquisition is halted if a fault is detected in the imaging. (**H**) Screenshot of acquisition pipeline software. Colored tiles indicate imaged stacks in different stages of processing.

**Figure S2.**
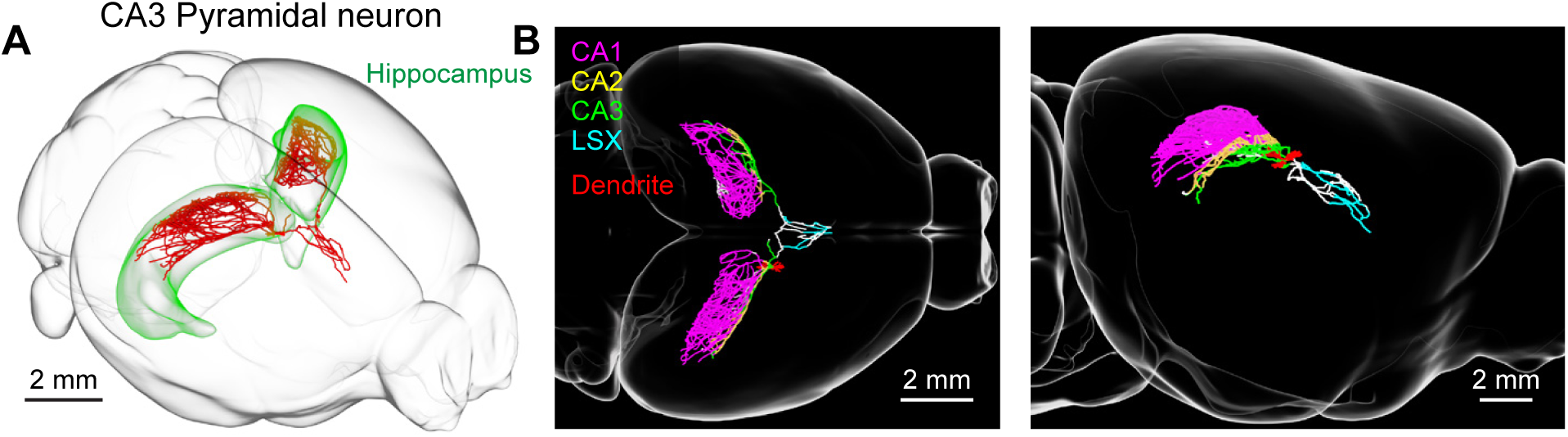
Reconstructed CA3 pyramidal neuron. Related to Figure 2. (**A**) 3d visualization of reconstructed pyramidal neuron (red) in CA3 of the hippocampus (green). (**B**) Left, horizontal view of the same neuron with axon color coded by its position in CA1 (magenta), CA2 (yellow), CA3 (green), or lateral septal complex (LSX; cyan). Dendrite is shown in red. Right, sagittal view of same neuron.

**Figure S3.**
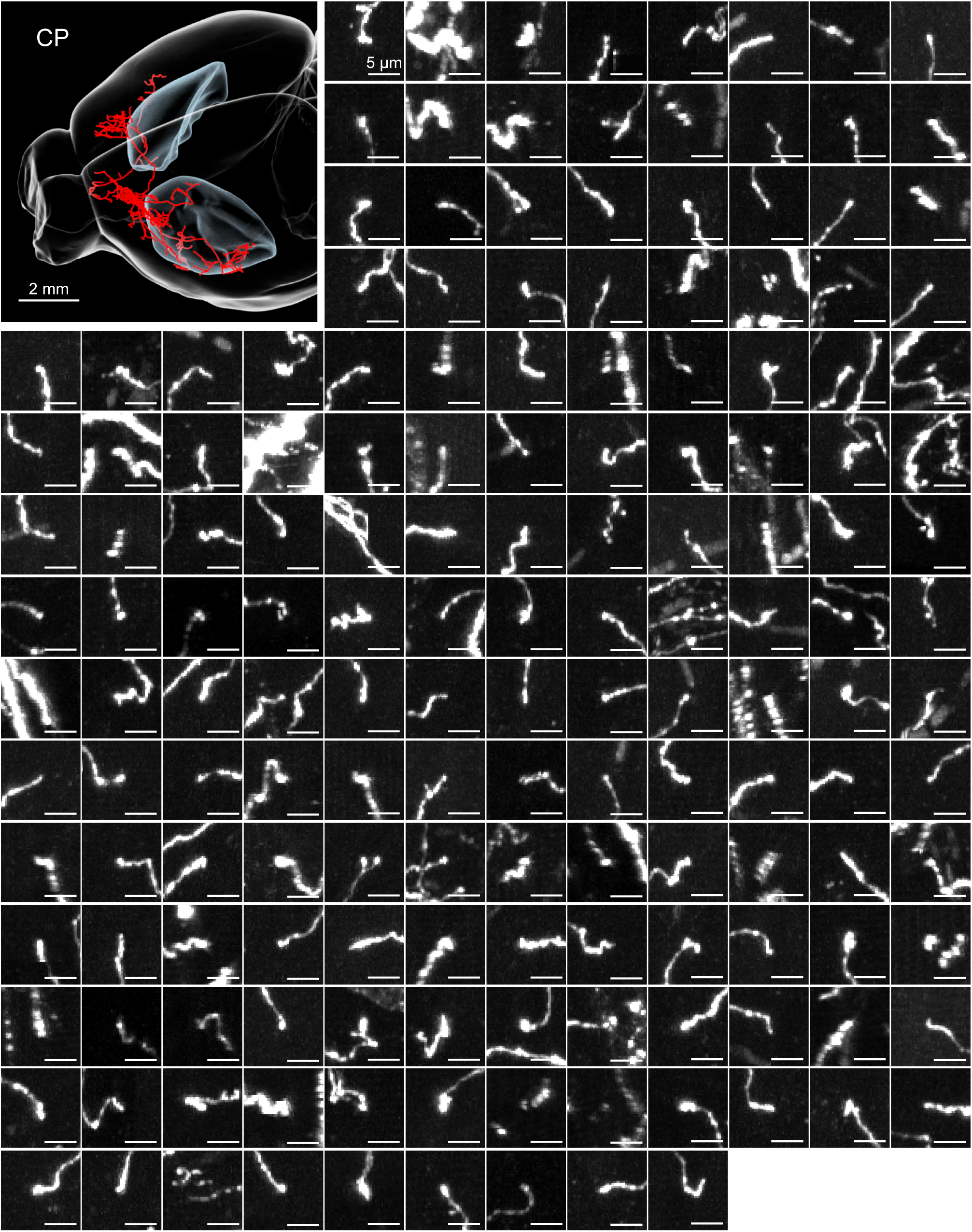
Axonal endings of a single motor cortex neuron. Related to Figure 2. Top-left, 3d visualization of a intratelencephalic neuron in the motor cortex. Caudoputamen (CP) is shown in blue. Other panels show maximum intensity projections of all identified axonal ends in the optimal viewing plane.

**Figure S4.**
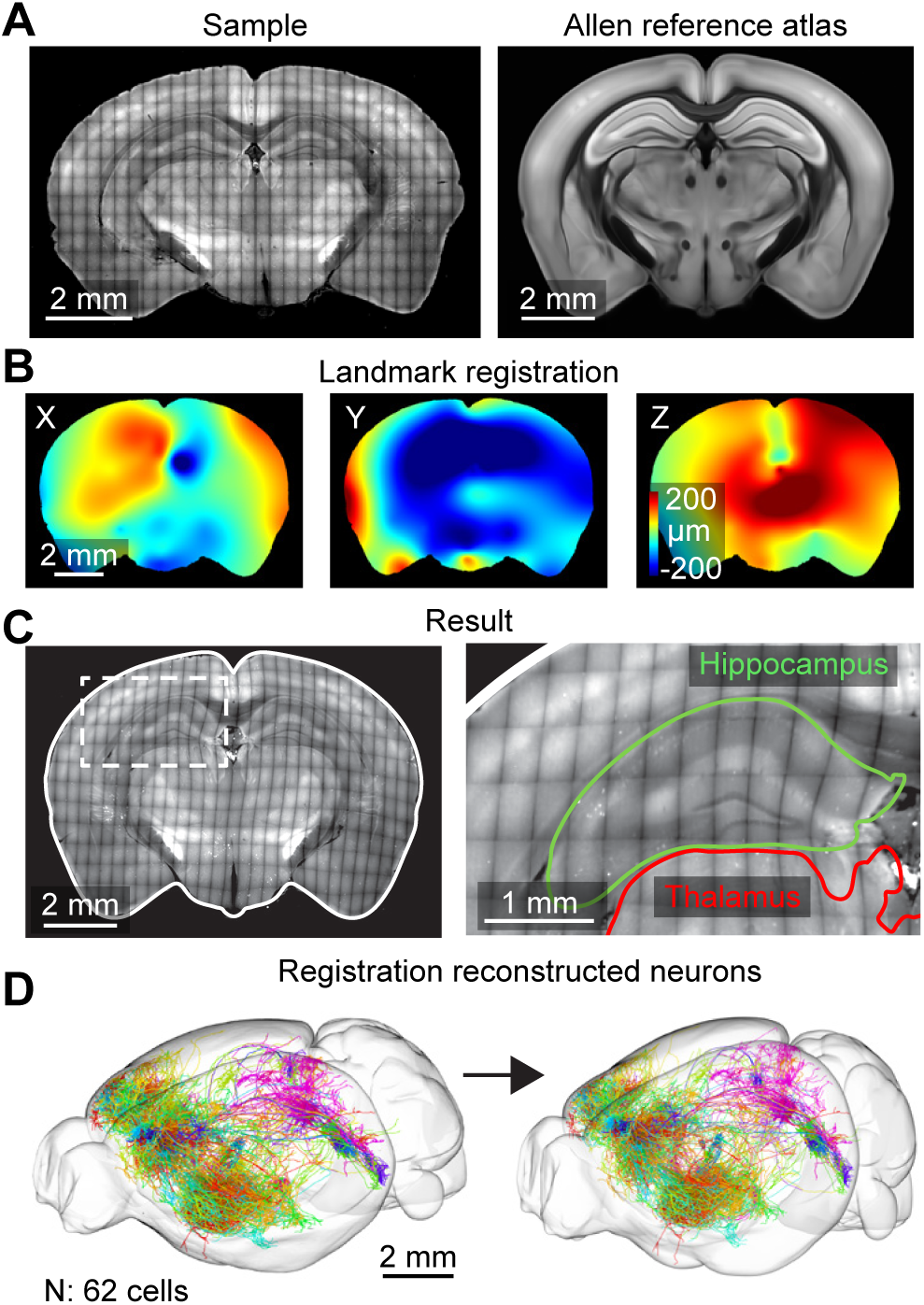
Registration of reconstructed neurons to the Allen Common Coordinate Framework. Related to Figure 2. (**A**) Coronal view of an imaged sample (left) and the corresponding section in the Allen reference atlas (right). (**B**) Spatial deformation of the imaged tissue in all three dimensions established by a landmark registration process. (**C**) Left, coronal section of same sample after registration. Outline of Allen reference atlas shown in white. Right, higher magnification view of area marked on left with an overlay of the boundary for the thalamus (red) and hippocampus (green). (**D**) Transformation of neuronal reconstructions to the Allen common coordinate framework. Left, 3d view of all reconstructions (n = 62 cells) within one sample. Right, same reconstructions after applying the transformation generated by the registration procedure.

**Figure S5.**
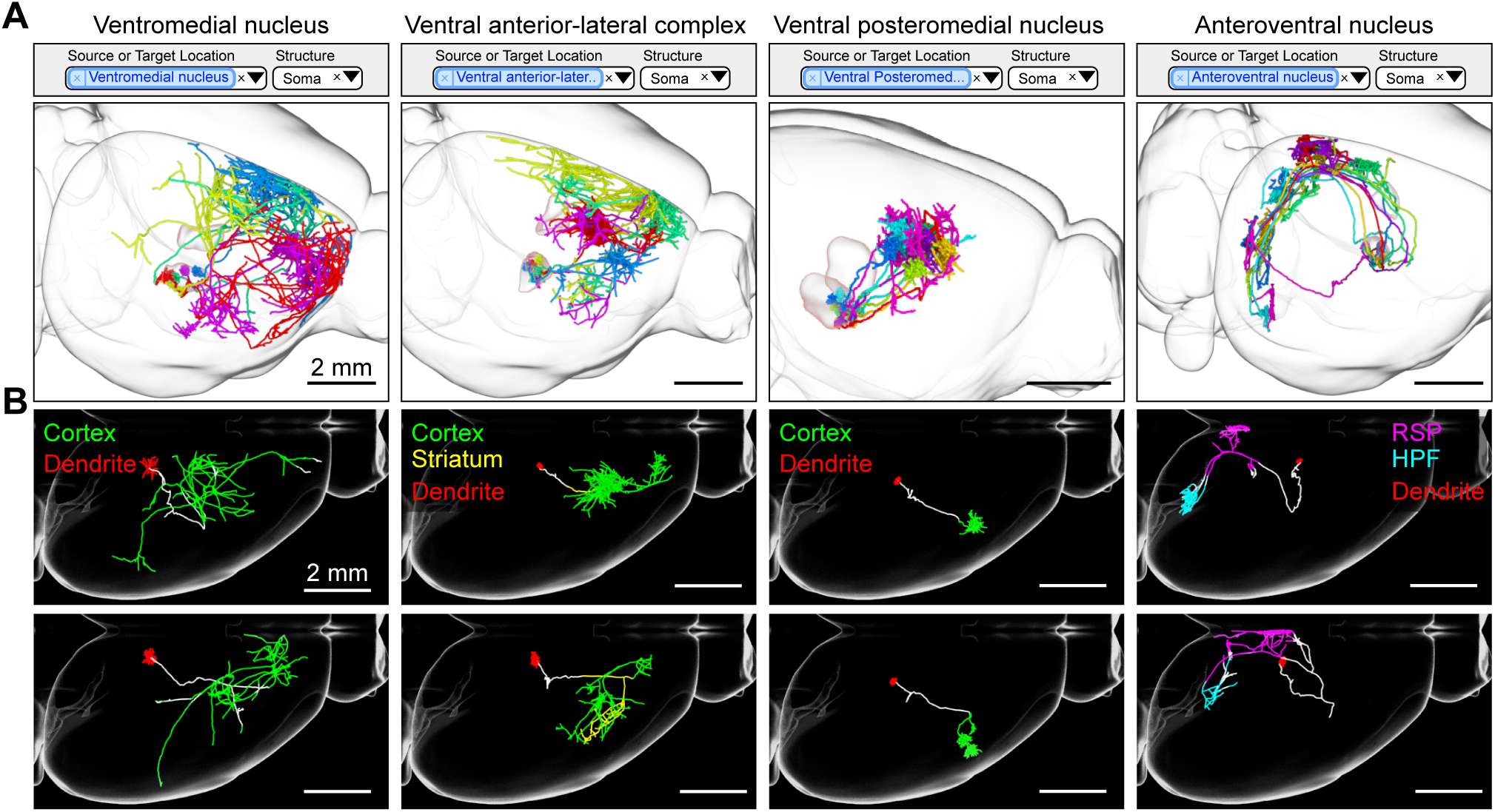
Projections from neurons across thalamic nuclei. Related to Figure 3. (**A**) Search queries (top) and 3d visualization (bottom) for neurons with somata in different parts of the thalamus: ventromedial nucleus, ventral anterior-lateral complex, ventral posteromedial nucleus, and anteroventral nucleus. (**B**) Example of single thalamic neurons in the same areas shown in A. Axons are color coded according to their anatomical position (RSP: retrosplenial cortex, HPF: hippocampal formation). Dendrites are shown in red.

**Figure S6.**
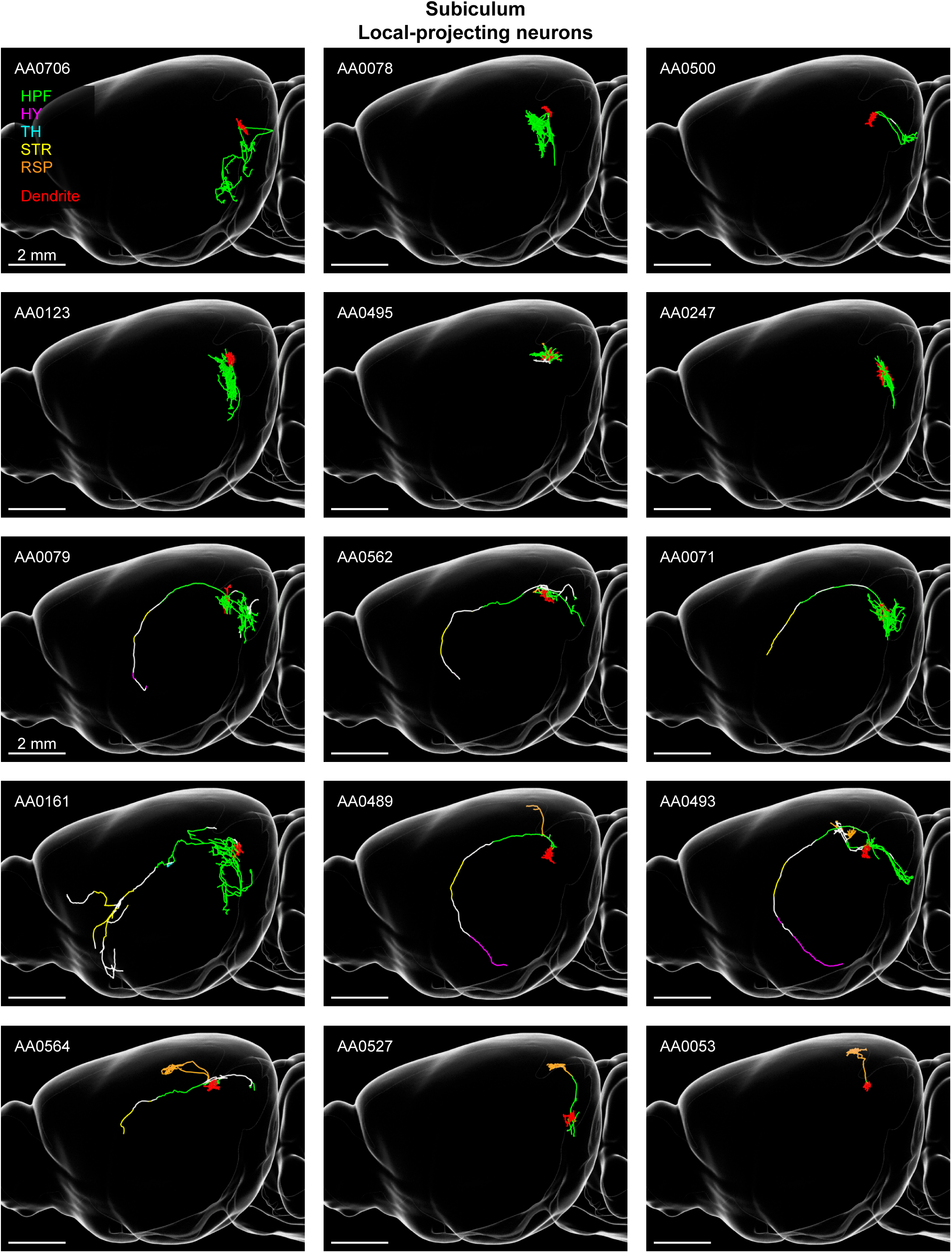
Local-projecting subiculum neurons. Related to Figure 4. Sagittal view of single subiculum neurons with local projections. Axon is color coded according to anatomical position (HPF: hippocampal formation, HY: hypothalamus, TH: thalamus, STR: striatum, RSP: retrosplenial cortex). Dendrite is shown in red. Unique neuron identifier in database shown in top-left.

**Figure S7.**
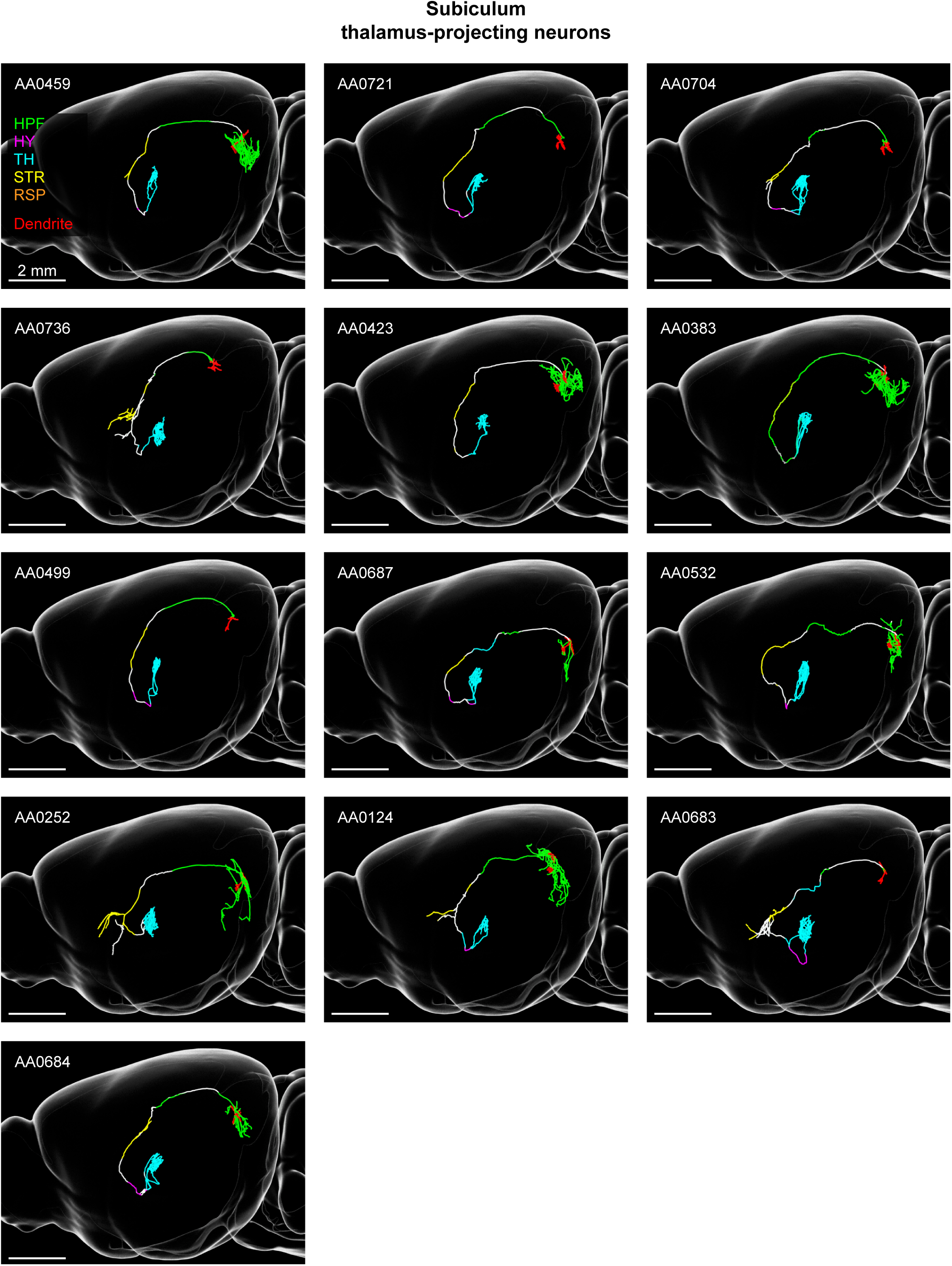
Thalamus-projecting subiculum neurons. Related to Figure 4. Sagittal view of single subiculum neurons with projections to the thalamus. Axon is color coded according to anatomical position (HPF: hippocampal formation, HY: hypothalamus, TH: thalamus, STR: striatum, RSP: retrosplenial cortex). Dendrite is shown in red. Unique neuron identifier in database shown in top-left.

**Figure S8.**
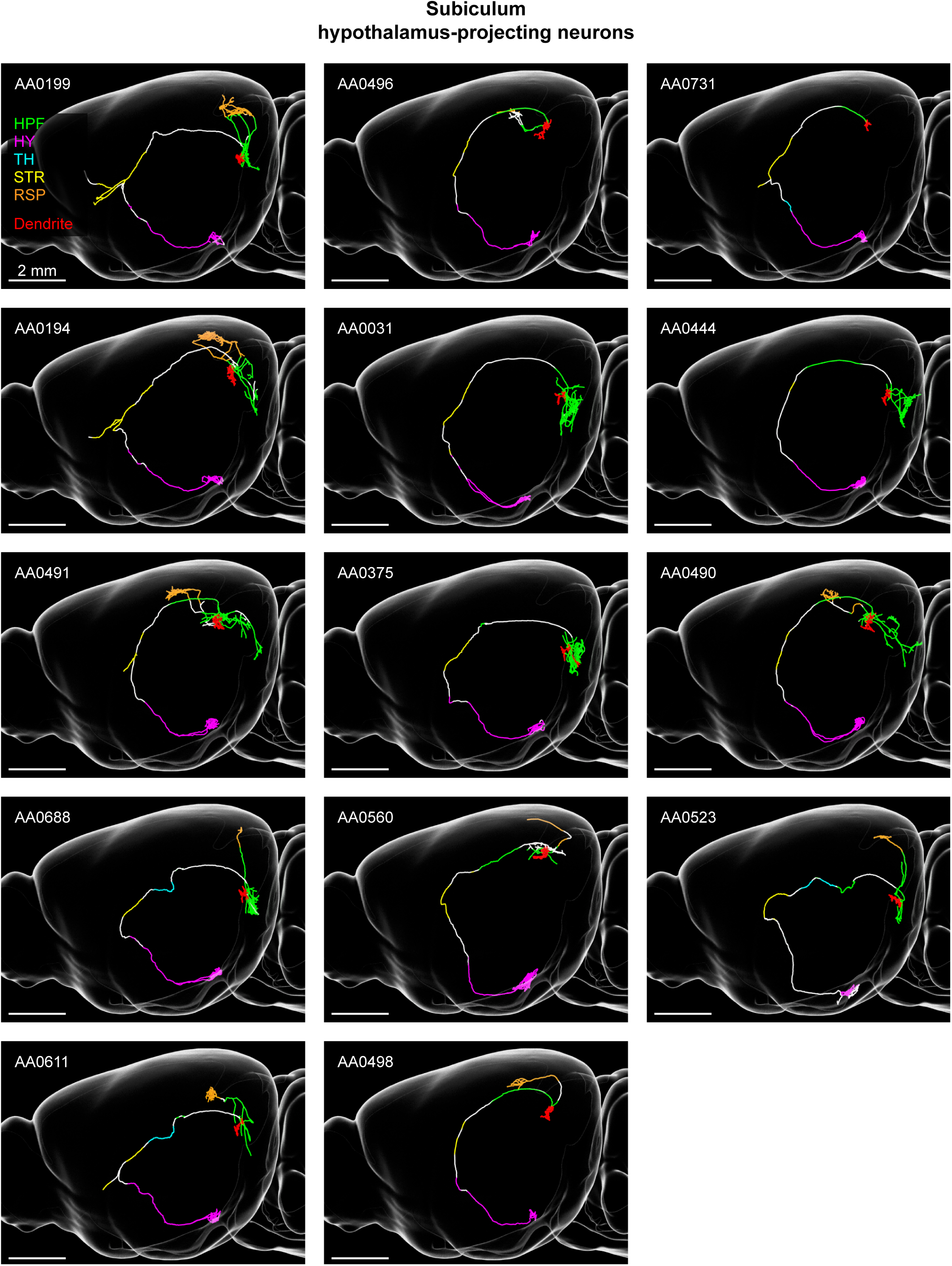
Hypothalamus-projecting subiculum neurons. Related to Figure 4. Sagittal view of single subiculum neurons with projections to the hypothalamus. Axon is color coded according to anatomical position (HPF: hippocampal formation, HY: hypothalamus, TH: thalamus, STR: striatum, RSP: retrosplenial cortex). Dendrite is shown in red. Unique neuron identifier in database shown in top-left.

**Figure S9.**
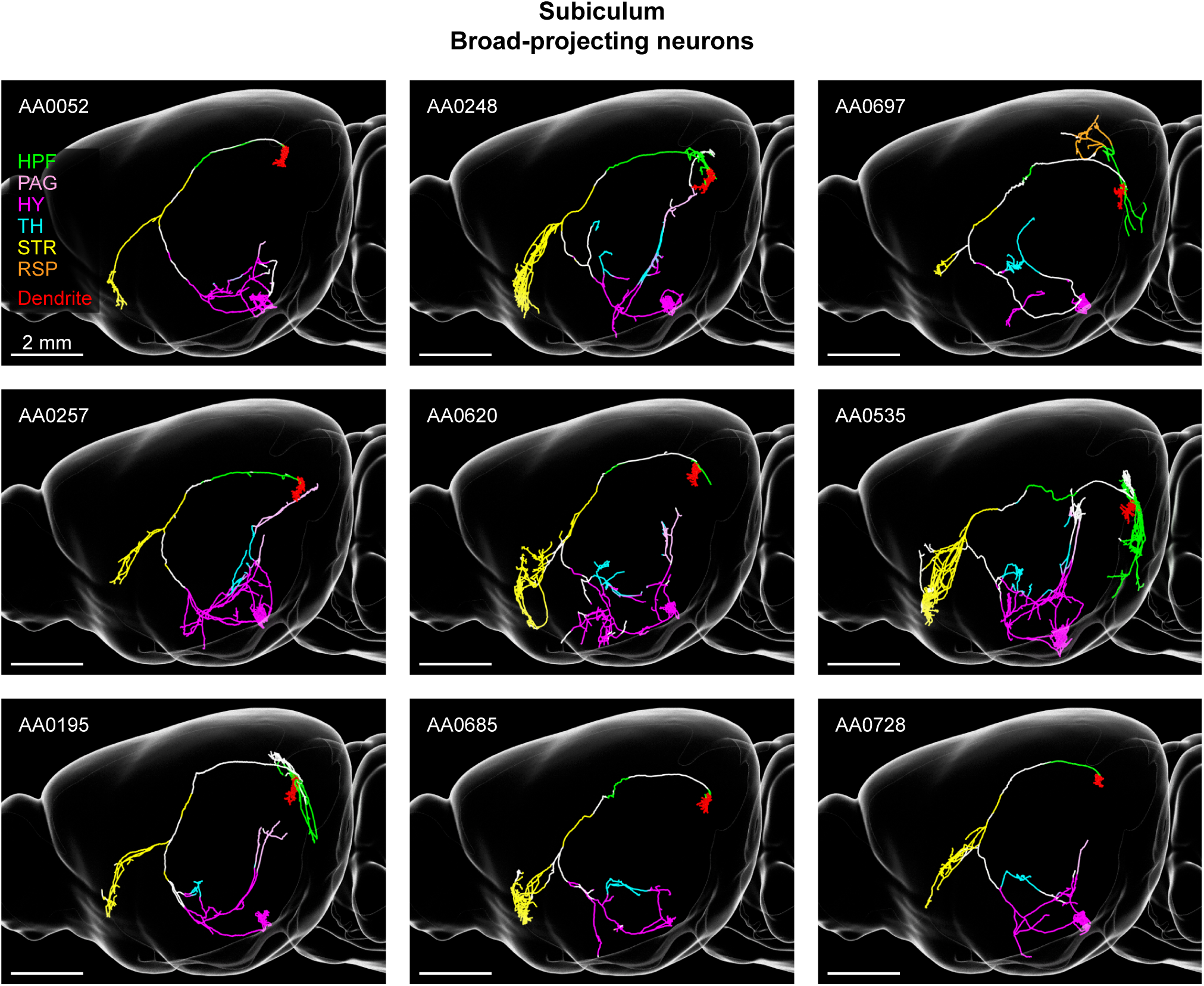
Broad-projecting subiculum neurons. Related to Figure 4. Sagittal view of single subiculum neurons with broad projections to multiple areas. Axon is color coded according to anatomical position (HPF: hippocampal formation, PAG: periaqueductal grey, HY: hypothalamus, TH: thalamus, STR: striatum, RSP: retrosplenial cortex). Dendrite is shown in red. Unique neuron identifier in database shown in top-left.

**Figure S10.**
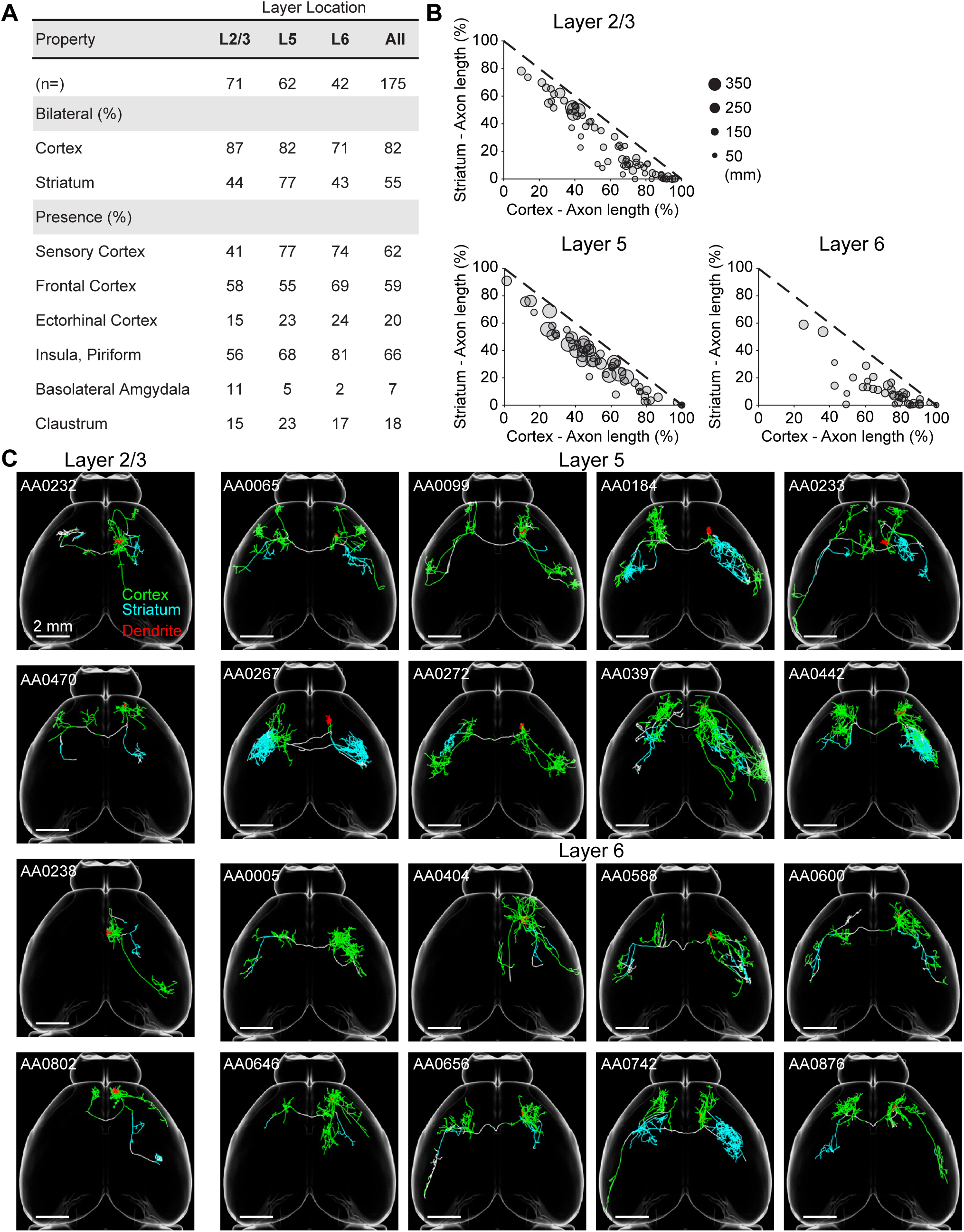
Projection patterns of intratelencephalic neurons in the motor cortex. Related to Figure 5. (**A**) Table of projection properties of intratelencephalic (IT) neurons in the motor cortex organized by laminar position. Only neurons with at least one axonal ending in a given area are counted. (**B**) Relationship between individual IT neurons axon length (as proportion of total length) in the cortex and striatum. Dotted line shows where combined length in cortex and striatum equals 100%. Neurons fall below the line due to axonal arbors in fiber tracts and other brain areas. (**C**) Horizontal view of individual IT neurons in different cortical layers. Axons are color coded according to their localization in the cortex (green) or striatum (cyan). Dendrites are shown in red. Unique neuron identifier in database shown in top-left.

**Figure S11.**
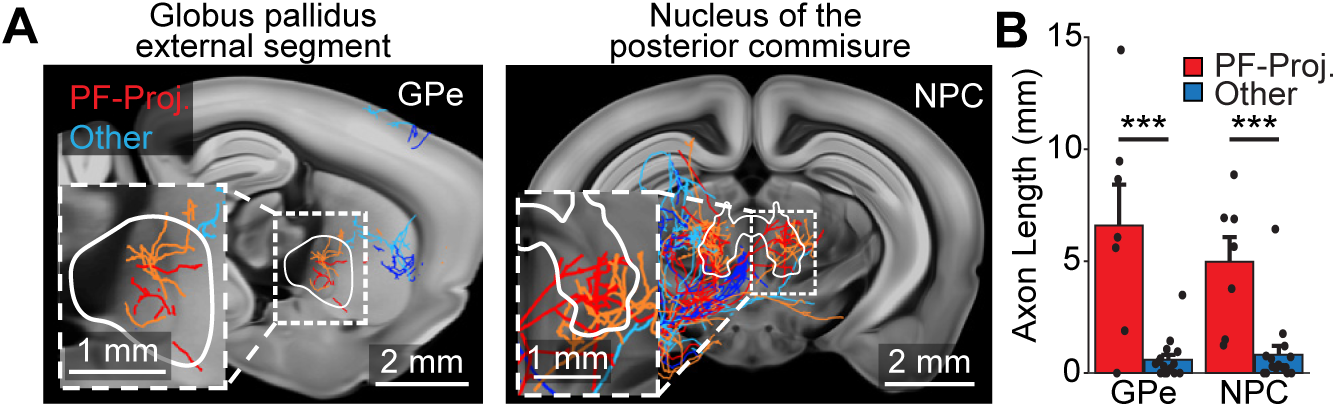
Specificity of PF-projecting pyramidal tract neurons. Related to Figure 6. (**A**) Left, sagittal view of PT thalamus-projecting neurons with arborizations in the parafascicular nucleus of the thalamus (red) and those without (blue). Outlined area denotes the globus pallidus, external segment (GPe). Right, coronal view of same neurons with outline of the nucleus of the posterior commisure (NPC). (**B**) Differences in axonal length between PF-projecting (red) and other PT thalamus-projecting neurons (blue) in the GPe and NPC. Error bars ± SEM. ٭٭٭p < 0.001, two-sample t-test. Greyscale images are from the Allen reference atlas.

**Figure S12.**
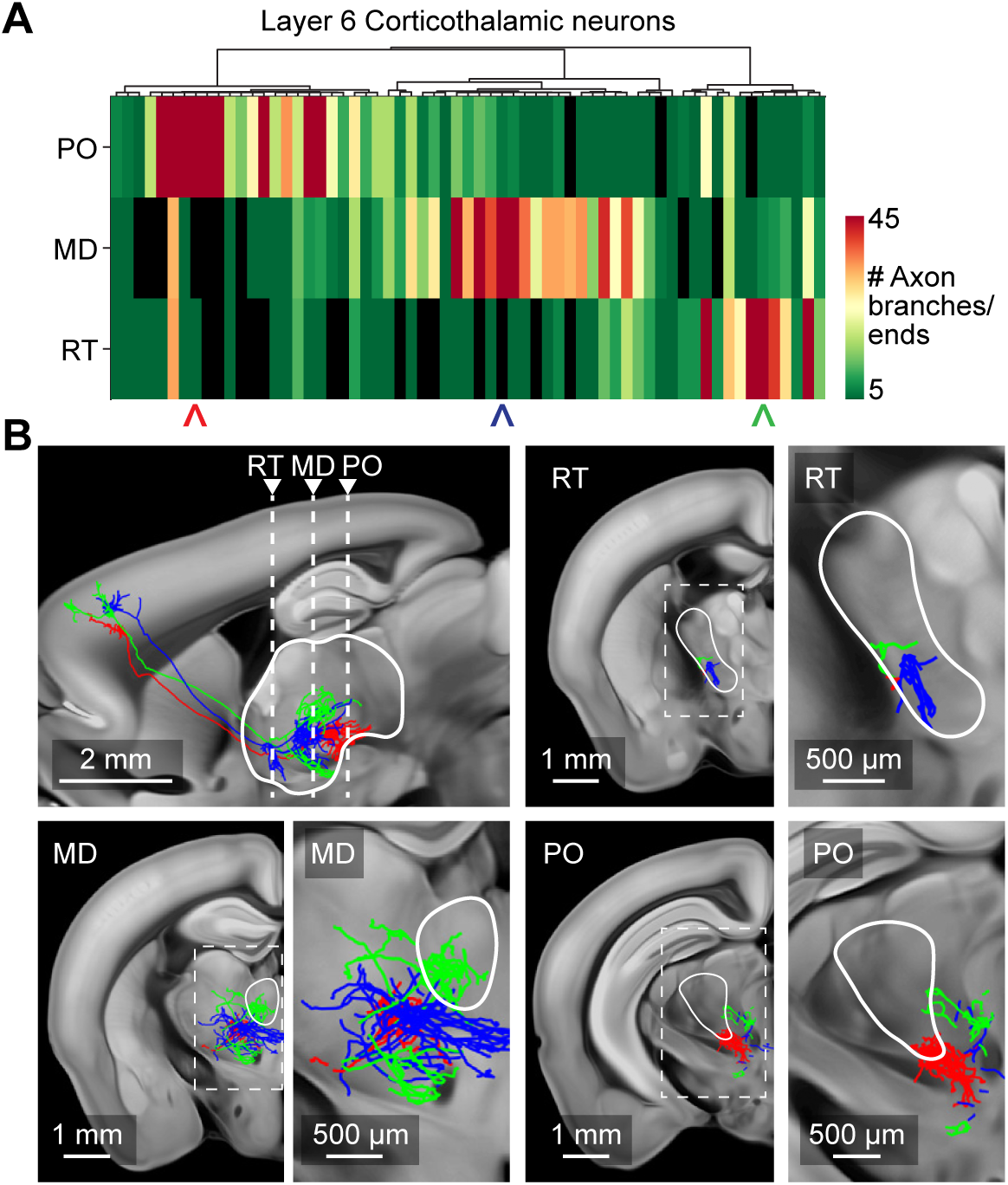
Thalamic innervation by layer 6 corticothalamic neurons in the motor cortex. Related to Figure 6. (**A**) Overview of thalamic innervation by layer 6 corticothalamic (L6-CT) neurons. Rows represent key innervated thalamic nuclei (PO: posterior complex, MD: mediodorsal nucleus, RT: reticular nucleus) and columns signify unique neurons. The color of the heat map denotes the number of axon branches and ends in that thalamic nuclei for a specific neuron. (**B**) Thalamic projections of example neurons highlighted in A (arrowheads). Top-left, location of shown coronal views marking the location of RT, MD, and PO.

**Figure S13.**
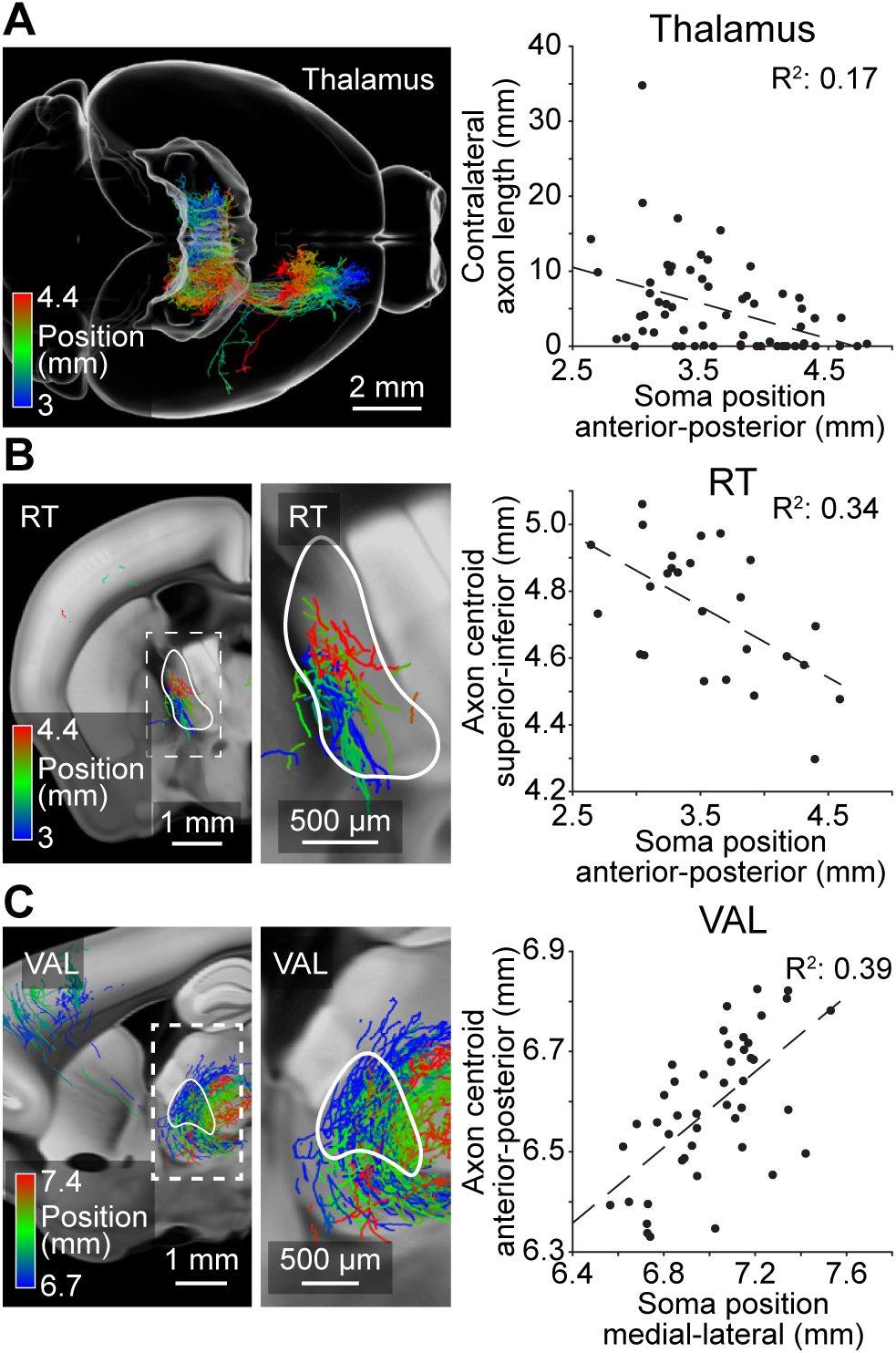
Topographic projections of layer 6 corticothalamic neurons in the motor cortex. Related to Figure 6. (**A**) Left, horizontal view of layer 6 corticothalamic (L6-CT) neurons in the motor cortex color coded by their anterior-posterior position. Right, relationship between somatic anterior-posterior position and axonal length in the contralateral thalamus. (**B**) Left, coronal view of neurons with projections in the reticular nucleus (RT) of the thalamus color coded by their anterior-posterior position. Inset, higher magnification view of dashed box on the left. Right, relationship between anterior-posterior position of the soma and axonal position in RT along the superior-inferior axis. (**C**) Left, sagittal view of axonal projections in VAL. Neurons are color coded by their medial-lateral position. Inset, higher magnification view of dashed box on the left. Right, relationship between somatic medial-lateral position and axonal anterior-posterior position in VAL. Trend lines depict linear fit. Axonal position of individual neurons is determined by the weighted centroid of their axonal projections within a specified region. Reported positions are in accordance with the Allen common coordinate framework. Greyscale images are from the Allen reference atlas.

